# Assessing climate change impacts for small-scale fisheries in the Gulf of California using Deep Learning

**DOI:** 10.1101/2025.03.28.645356

**Authors:** R. Cavieses-Nuñez, Q. Lu, H.N. Morzaria-Luna, P. Mallick, S. Kumara, C.R. Navarrete-Torices, Gabriela Cruz-Piñón, S. Buechler, K. Lopez-Olmedo

**Affiliations:** Instituto de Investigaciones Oceanológicas, UABC-CONAHCyT. Carretera Ensenada-Tijuana No. 3917, Fracc. Playitas, Ensenada, Baja California, C.P. 22860; Departamento Académico de Ingeniería en Pesquerías, Universidad Autónoma de Baja California Sur. Carretera al Sur Km 5.5, Apartado Postal 19-B, La Paz, Baja California Sur. México. C.P. 23080; Laboratory for Intelligent Systems and Analytics (LISA), Harold and Marcus Inge Department of Industrial and Manufacturing Engineering, The Pennsylvania State University, 233 Leonhard Building, University Park, PA 16802; CEDO Intercultural. Puerto Peñasco, Sonora, Mexico 83550 & Tucson, AZ. USA 85733; Visiting Researcher, Northwest Fisheries Science Center, NOAA. 2725 Montlake Blvd. Seattle WA 98112, USA; Departamento Académico de Ciencias Marinas y Costeras. Universidad Autónoma de Baja California Sur. Boulevard Forjadores S/N, La Paz, Baja California Sur. México. C.P. 23080; Associate Research Professor. Women’s Engagement with Agricultural Research and Education (WEARE) program and International and National Extension. Affiliation: Agricultural Sciences Global at The Pennsylvania State University

**Keywords:** climate change scenarios, forecast, fishing communities, artificial intelligence, deep learning

## Abstract

Small-scale, multispecific fisheries in the Gulf of California face significant challenges including limited species-specific catch data, uncertainty about climate change impacts, and insufficient biological information needed for traditional deterministic models. These knowledge gaps hamper efforts to forecast future conditions and develop appropriate management strategies accurately. The complexity of these multi-species fisheries, combined with data scarcity for many target species, creates substantial barriers to quantifying and addressing climate vulnerability. Deep learning approaches offer a promising alternative by leveraging available data to identify patterns and project trends despite these limitations, providing valuable insights for fisheries management in data-poor contexts. Here, we apply a Mixture of Expert, a deep learning forecasting models for small-scale, multi-specific fisheries in the Gulf of California under future climate change scenarios. Results show varied responses across marine habitats, with reef and benthic fish projected to experience substantial declines (-12.46% and -9.37%) during the 2050s-2060s, followed by recovery in the 2070s-2080s. Economic implications are significant, with reef fish facing projected losses of $1.2 million by the 2050s before recovering by the 2080s. Shapley Additive Explanations (SHAP) analysis was applied to evaluate the importance of features for each predictive model, the analysis revealed the effects of the temperature in different depths for each fishery, and the sensitive analysis pointed to the magnitude of the effect. Our findings suggest that climate impacts will not be uniform across the Gulf, necessitating region-specific management approaches and highlighting the value of maintaining diverse fishing portfolios to enhance resilience against climate-driven changes.

## Introduction

Climate change threatens marine ecosystems and human well-being through changes in temperature, oxygen, and other biogeochemical characteristics [1]. As a result, multiple impacts on ocean systems are expected, including alterations in species distribution, growth, reproduction, abundance, and trophic relationships [2]. Due to this, the effects on global fisheries will be of two general types: changes in fishery productivity, which will affect potential yields and profits, and changes in species distributions, which will affect the spatial and temporal distribution of fishing efforts, the species targeted, the location where fish are landed and processed, and the characteristics of fishing fleets [3,4], which may influence food web function [5] and lead to further unexpected effects for commercial species such as population collapse [6] or regime shifts [7].

Overall, climate change will affect the environmental stochasticity of marine ecosystems, affecting fishery stocks and profits, disrupting the value chain, and significantly affecting food security and the way of life for fishing communities [2,8]. Globally, fisheries production has already decreased due to ocean warming, with some regions experiencing up to 35% declines in the maximum sustainable yield (MSY) of important fish stocks since the 1930s [9]. Under a future climate change high-emission scenario (RCP 8.5) [10], global fish biomass may further decrease by 30-40% in tropical regions by 2100 [11]. Effective and adaptive fisheries management frameworks are critical to limit or mitigate the impacts of climate change on fisheries [2].

Rapidly changing ocean conditions under climate change are a challenge for small-scale fisheries, which provide food and nutrition security, contribute to health and well-being, and provide income in areas that often lack other economic opportunities [12]. Globally, 60 million people are employed along the small-scale fisheries value chain, and almost 500 million depend partially on small-scale fisheries [13]. These fishers use their knowledge and personal experience to maximize catches by predicting the behavior and movements of target species. Small-scale fisheries’ catches show marked seasonality and spatial variation [14] and they are highly vulnerable to shifts in target species’ spatial distribution, phenology, and seasonality, which are all predicted to be impacted by climate change [15].

From a cybernetics perspective, which studies control and information processes in complex systems [16,17], small-scale fisheries can be understood as information-processing networks where each component - environment, ecosystem, and fishing community - continuously exchanges and processes signals that modify the behavior of the system as a whole [18,19]. Small-scale fishers constantly adapt by interpreting signals from the environment (such as temperature and productivity), the resource (abundance and distribution), the market (prices and demand), and the regulatory framework, developing specialized knowledge about their target species [20,21]. This information-processing network, coupled with the scarcity of detailed biological data, requires approaches that emulate both the control processes and expert knowledge within the system. Within this context, there is an urgent need for robust projections of the effects of climate change on fisheries, including future catch trends, that could help manage resources and the market by identifying species currently present occasionally but will become dominant [14,22].

The synchronous relationship between fisheries catch and ocean conditions allows for the development of catch forecasts [23]. However, this relationship is complex, non-linear, and time-lagged [24,25]. Ocean physical variables are non-linear; in particular, ocean temperature is high-dimensional and difficult to model mechanistically; causal explanations are hard to generate, and meanwhile, biological populations are not just tracking the environment, but rather, ecological dynamics are characterized by nonlinear amplification of stochastic physical forcing [4,26]. Constructing a mechanistic model that can account for these complex relationships is a challenging task [27]; however, recent breakthroughs in integrated climate–fishery models that can learn directly from the data have greatly enhanced our ability to forecast how future climate scenarios might affect catch volumes [28–32]. Deep Learning models can learn the stochastic dependency between the past and the future and the nonlinear relationship between the variables [33]. However, catch forecasting is limited by data availability; therefore, developing approaches that predict future catches with little information and effort will be critical to fisheries’ management [28].

In the Gulf of California, Mexico, small-scale multispecific fisheries, involving more than 80 target species of fish, sharks, and invertebrates [34], are vital for food security and local economies [35–37]. Climate change impacts are expected to change the spatial and/or temporal distribution of fishing effort for small scale fisheries in the Gulf of California [38], the species targeted, the location where fish are landed and processed, and/or the characteristics of fishing fleets [38], and resulting in a decrease in catches may significantly affect food security and the way of life in these fishing communities [39]. These small-scale fisheries are also susceptible to non-climatic drivers such as market changes, demographics, and overexploitation [40]. Therefore, assessing fisheries’ vulnerability to climate change and developing adaptive management responses for small-scale fisheries in Mexico is crucial [41]. However, Mexico’s fisheries management system operates with insufficient biological data on fishery stocks, including species diversity, distribution, life history, and catch spatial patterns necessary to develop deterministic models to predict climate change impacts on all fisheries [42].

We apply deep learning models to construct forecasting models for small-scale, multi-specific fisheries in the Gulf of California under future climate change scenarios. We seek to provide comprehensive insights into climate change’s potential impacts on small-scale fisheries’ catch dynamics and contribute to informed management strategies for sustainable resource utilization in the region. We aim to address the following main questions: (a) Can deep learning models demonstrate sufficient predictive efficacy to anticipate catch volumes for these fisheries amidst evolving future ocean warming?, (b) how much will fisheries catch be affected under future warming scenarios, and c) what are the implications of catch changes for fishers and fishing communities? It is important to note that our approach does not provide specific reference points for fisheries management (i.e. catch quotas). Rather, our models are a tool to understand and identify patterns in potential catch dynamics and project trends that can help stakeholders anticipate and adapt to future changes in the fishery system.

## Methods

We apply deep learning models that relate catch data to ocean temperature to forecast changes under future warming scenarios. We use catch landing reports from throughout the Gulf of California, which are categorized into fishery regions and relate them to potential seawater temperature. Our methodology follows several key steps: (1) georeferencing catch landing reports to assign ocean temperature values, (2) conducting cluster analysis to derive eight distinct fishery regions, (3) categorizing species by habitat groups, (4) obtaining and downscaling sea temperature data from GLORYS and MPI-ESM-MR climate projections, (5) developing a Mixture of Experts (MoE) model that combines LSTM and CNN architectures to capture the complex non-linear relationships between environment and catch, (6) validating model performance using multiple metrics (R2, MSE, accuracy), and (7) conducting sensitivity analyses using SHAP values to understand feature importance. We apply this integrated framework to forecast catch under future warming scenarios through 2085, focusing on both ecological patterns and economic implications.

### Gulf of California

The Gulf of California borders six Mexican states: Baja California, Baja California Sur, Sonora, Sinaloa, Nayarit, and Jalisco [43]; it is Mexico’s marine ecosystem with the highest fishery production [44,45]. The Gulf’s rich biodiversity sustains unique marine species, emphasizing the importance of sustainable fishing practices for ecosystem health and global food supply [46,47]. The Pacific Ocean, including the Gulf of California, yields over 633 thousand tons of fish and shellfish annually, contributing 57% of Mexico’s fish production [48]. Gulf of California’s fisheries are integrated into regional diets and supply food and income for local communities; they also provide a significant source of seafood for consumption throughout Mexico and for import to the United States [49,50]. Small-scale fisheries in the Northern Gulf target over 80 fish species, sharks, and invertebrates [34]; they share fishing areas, fishing seasons, fishing gear, and fleets [32]; and are important culturally, economically, and for food security [34].

The Gulf of California’s marine ecosystems are already experiencing the impact of climate change. Temperature measurements show that the Gulf’s waters have been warming steadily, with a documented annual temperature increase of 0.04°C from 1998-2015 [51]. This thermal change manifests in extended summer periods and broader warm water regions throughout the Gulf [35]. Increased wave intensity has been observed along the coastline, with resulting erosion affecting approximately two-thirds of the Gulf’s shores [52]. Furthermore, projections suggest that 2050 warming waters will substantially reduce populations of commercially valuable reef fish species in areas below 25°N latitude, potentially altering reef ecosystem dynamics [53–55].

### Fisheries data

The main source for fishery data in Mexico are catch landing reports submitted to the National Fisheries Commission (CONAPESCA), online and through local fishing offices fishery permit-holders (https://conapesca.gob.mx/wb/cona/avisos_arribo_cosecha_produccion). The catch landing reports contain information about monthly catch, prices, and landing zones, and they commonly aggregate catch from different fishing trips and economic units (vessels) operating under the same permit.

We used catch landing reports for the main species targeted by small-scale fisheries [56,57] between 2002 and 2023, due to the disponibility of data from CONAPESCA. We extracted the following fields from each catch report: fishery office, state, date, permit number, fishing zone, landing site, common species name, weight of fish landed, and the catch price. The landing reports were corrected for misspellings and consistency in the common species names and landing site names.

Although the catch landing reports include the names of fishing zones and landing areas, they do not include geographic coordinates. Since we needed to assign each catch landing report to a specific geographic location to assign an ocean temperature value, we used one of the following approaches to georeference the landing records: 1) We cross-referenced landing sites with the Atlas of Fisher Locations for the Mexican Pacific (available from: https://www.conapesca.gob.mx/wb/cona/atlas_de_localidades_pesqueras), 2) We searched for landing sites’ names in the Catalog of Geostatistical State and Municipal Areas and Localities, developed by the National Institute for Geography and Statistics (INEGI), available here: https://www.inegi.org.mx/servicios/catalogounico.html, 3) We searched for landing sites using the Google Maps Platform Place Search API. We reviewed georeferenced points for name consistency, 4) We used the address of the permit holder obtained from the National Registry for Fisheries and Aquaculture (RNPA) requested through the National Institute for Transparency, Data Access and Protection of Personal Information https://home.inai.org.mx/. As a result, each catch landing report was associated with a geographic point along the coastline or further inland (Figure 1). To assign ocean temperature values to a given catch landing report, we first assigned them to fishery regions (below).

**Figure 1.**
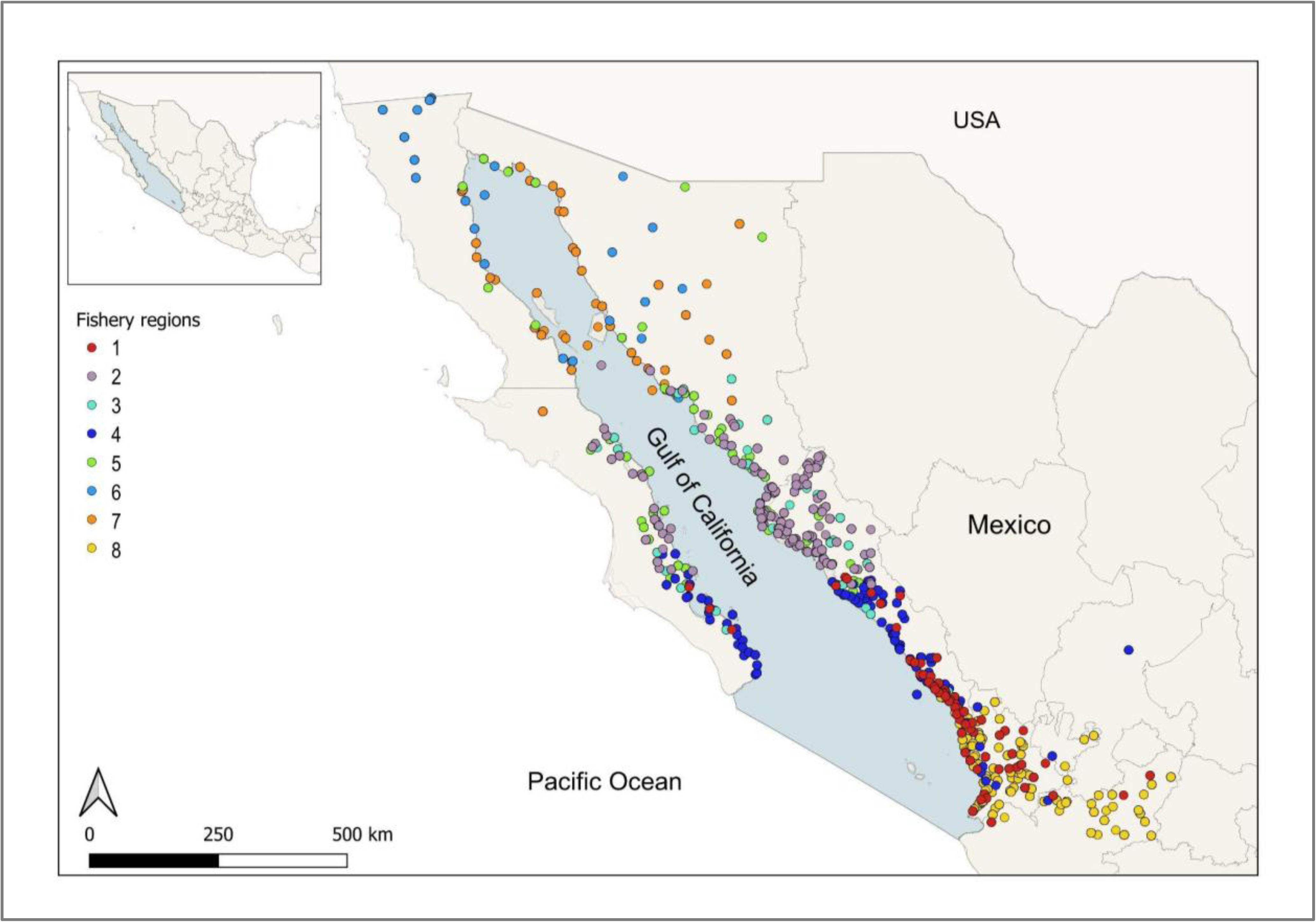
Catch landing records by fishery regions. Points indicate catch landing records assigned to landing sites or permit holder address; see text for detailed explanation.

### Fishery regions

We conducted a cluster analysis on the georeferenced catch landing reports to derive eight fishery regions, using the latitude, temperature, and species caught as explanatory variables. For the fisheries within each region, we created two distinct time series: (1) monthly catch for each species, aggregated by summing the catch of all economic units by month, and (2) monthly catch where data from each economic unit remained unaggregated, with only the time steps consolidated on a monthly basis. The time series were arranged in a pandas data frame with the following columns: date, fishery region, species name, and catch weight. To simplify the analysis of results, we grouped the fisheries by their typical habitat (Table 1.)

**Table 1.** Catch categories by habitat group. Catch species are the local original common name; see Supplementary Information S1 for species Spanish and scientific names. Fish habitats were taken from FishBase [58].

| Habitat Group | Catch species |
| --- | --- |
| Benthic invertebrates | clam, shrimp, blue crab, snail, lobster, octopus, sea urchin, oyster, sea cucumber, abalone |
| Pelagic fish | tuna, skipjack, bonito, jack, pompano, sierra, sardine, shark |
| Demersal fish | croaker, grouper, baqueta, corvina |
| Benthopelagic fish | bass, grunt, mullet, mojarra, bandera |
| Reef fish | cabrilla, red snapper, amberjack, snapper, pierna, lane snapper, villajaiba, snook |
| Benthic fish | flounder, rays |

### Ocean temperature data

As our primary environmental variable, we used potential sea water temperature, which refers to the temperature that a defined parcel of water would have if it were moved adiabatically to a standard pressure level. In this context, a “parcel” of water is a small, hypothetical volume used to study the properties and behavior of water in the ocean. Potential sea water temperature provides a more stable metric for analyzing temperature changes across different depths and regions. We obtained sea water potential temperature from the Global Ocean Physics Reanalysis (GLORYS) developed by the E.U. Copernicus Marine Service Information [59] (https://doi.org/10.48670/moi-00021). GLORYS is an eddy-resolving global ocean reanalysis (1/12° horizontal resolution, 50 vertical levels) covering the altimetry from 1993 onwards. We used seawater potential temperature at a 0.083° × 0.083° resolution for 0-500 m depth from December 1993 to July 2024. We downloaded GLORYS data files subsetted to our study region using the Copernicus Marine Toolbox API in Python (https://help.marine.copernicus.eu/en/articles/7972861-copernicus-marine-toolbox-cli-subset).

### Future ocean warming

The future ocean temperature was derived by empirically downscaling global ocean climate projections based on the GLORYS model grid following [60]. We used an empirical delta-downscaling approach to project projected step changes (i.e., annual changes 1993-2024) in temperature from a coarse-scale Earth System Model to the higher-resolution climatology from GLORYS to derive a time series of temperature from 2002 to 2023, corresponding to the catch data, and forward 50 years to 2085.

We used global climate projections from the MPI-ESM-MR Earth System Model developed by the Max-Planck-Institut für Meteorologie [61] as part of the CMIP6 (Coupled Model Intercomparison Project 6), which contributed to the 6th IPCC report [62]. We used future emissions projections with social factors [63], the Shared Socioeconomic Pathways (SSPs), which detail scenarios that explore interactions between climate change and global socioeconomic futures [64] and outline potential societal and ecosystem development trends during the 21st century [65]. We focused on the SSP5 (ssp585), indicative of fossil fuel-driven development, and SSP1 (ssp126), representing sustainability, for the 2050 horizon. The future emission projections part of the SSPs are the Representative Concentration Pathways, which outline future physical impacts of climate change based on radiative forcing levels at 2100 in the tropopause, compared to pre-industrial levels [66,67] (refer to Supplementary Information S2).

### Deep learning models

The modeling process is based on two key assumptions. First, landing reports are assumed to exhibit a continuous and consistent rate of uncertainty across all data points, ensuring that potential biases are evenly distributed. Second, an indirect and nonlinear relationship is presumed between fishing catches and sea temperature, reflecting the complex interactions between environmental conditions and fishery dynamics. These assumptions guide the model’s structure and interpretation, providing a foundation for its predictive capabilities.

This study employed a deep learning approach involving a Mixture of Experts (MoE) model, Convolutional Neural Networks (CNNs), and Long Short-Term Memory (LSTM) networks. MoE can emulate the adaptive nature of fisheries by combining different specialized neural networks, each processing specific aspects of the system, analogous to how different actors process information in the real fishing system. MoE includes a gating network that learns to assign each data instance to the most appropriate expert [69,71]. Although no direct applications of MoE in fisheries exist, this method is well-suited for complex datasets such as fisheries data, which often exhibit both temporal and spatial heterogeneity because it can simulate adaptive processes in complex systems [68,69], making it highly relevant for fisheries modeling. By allowing different experts to handle distinct portions of the input space, the MoE can effectively model systems characterized by multiple regimes or patterns [72,73]. The gating mechanism then optimally integrates the output of these experts. Such specialization is particularly valuable in the Gulf of California’s multi-species fisheries, where species often respond differently to shifting ocean conditions. Moreover, the MoE framework balances model complexity with interpretability, thereby improving predictive accuracy [72].

### Computational requirements

We used the Google Cloud Computing framework, with a virtual instance with 32 cores and 124 GB RAM. The virtual machine used Linux, and a Conda environment using Python 3.14. We developed the models using pandas, Numpy, Keras, and TensorFlow libraries. The input data had a tabular shape with 689,399 rows x 33 columns, where rows are individual reports and the columns used where: Id, date, locality name, state name, fishery office, fisheries species name, landed weight in kg, value in Mexican pesos, region label, decimal longitude, decimal latitude, mean temperature at 30 m, mean temperature at 10 m, and temperatures from 6 m, 8 m, 9.5 m, 11.4 m, 13.5 m, 15.8 m, 18.5 m, 21.6 m, 25.2 m and 30 m, some others columns where dropped by duplicity or because they were not relevant for the study.

### Initial 2002-2023 time series model development

#### Preprocessing

The data frame incorporates both fisheries and environmental variables, which are categorical (such as species and office names) and numerical (SST, cats, values, and frequency). Categorical data was binary encoded, and numerical data was standardized using a min-max scaler. We then divided the data frame into training and validation sets, comprising 80% and 20% of the data. The forecast target was set as the one-step future catch weight, with all DataFrame variables serving as inputs.

### Model development with data for 2018-2023

As a first step, we examined various deep neural networks like Long-Short Term Memory (LSTM) and Convolutional Neural Networks (CNN) architectures to predict the target variables. LSTM neural networks were tested due it’s ability to find patterns and forecast time series [70]. The CNN were used to test the capability to find spatial patterns, in this case we also tested one dimension CNN model [71].

We employed Tree-structured Parzen Estimators (TPE) [72], a Bayesian optimization method, to tune hyperparameters of our deep neural network. Implemented through the Optuna framework, TPE guided the selection process based on prior evaluations to efficiently identify optimal network configurations, including layer depths, neuron counts, and learning rates. This approach significantly streamlined the Mixture of Experts (MoE) model’s development cycle by focusing computational efforts on more promising regions of the hyperparameter space.

### General model “First generation”

The first MoE model was trained using aggregated data, where monthly catches for each species within each fisheries region were summed to produce a single value per month. While this approach simplified the temporal structure of the data, it significantly reduced the number of training examples available to the model, limiting its exposure to variability and potentially affecting its generalization capability. Despite these constraints, this “first generation” model provided a baseline for understanding the temporal patterns in aggregated monthly catch data.

### Expert model “Retrained Model”

The model was retrained to address the limitations of the aggregated data approach, which uses disaggregated data from arrival notices. Instead of monthly sums, this dataset preserved individual arrival records, effectively increasing the number of examples available to the model and enhancing its capacity to capture fine-grained patterns and variability in the data. The increase in training samples provided a richer dataset for the model to learn from, allowing for a more robust representation of the underlying dynamics within each fisheries region and species. The structure of the model can be seen in Table 2.

**Table 2.**
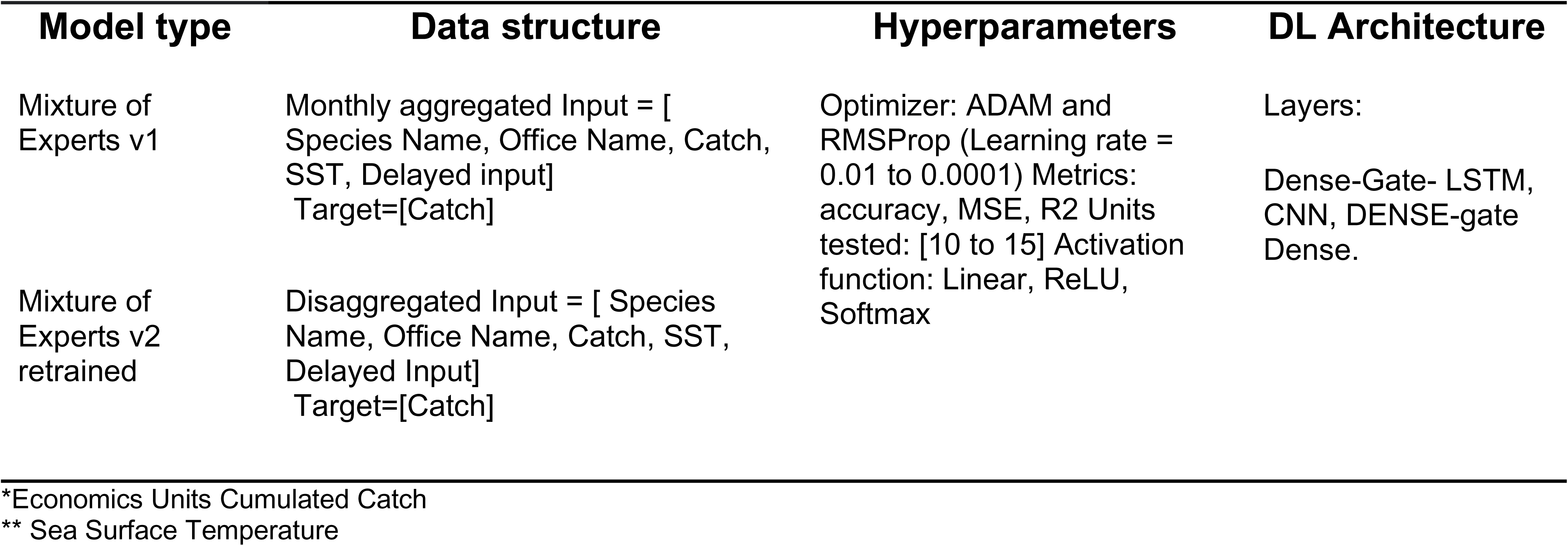
Structure of the tested model.

| Model type | Data structure | Hyperparameters | DL Architecture |
| --- | --- | --- | --- |
| Mixture of Experts v1 | Monthly aggregated Input = [ Species Name, Office Name, Catch, SST, Delayed input]<br>Target=[Catch] | Optimizer: ADAM and RMSProp (Learning rate = 0.01 to 0.0001) Metrics: accuracy, MSE, R2 Units tested: [10 to 15] Activation function: Linear, ReLU, Softmax | Layers:<br>Dense-Gate- LSTM, CNN, DENSE-gate Dense. |
| Mixture of Experts v2 retrained | Disaggregated Input = [ Species Name, Office Name, Catch, SST, Delayed Input]<br>Target=[Catch] |  |  |

\*Economics Units Cumulated Catch
\*\* Sea Surface Temperature

### Validation process

We applied data segmentation to validate the model’s performance, creating an 80% training process and using 20% of the data for validation. We used diverse performance metrics to evaluate the models, such as accuracy, mean squared error (MSE), correlation coefficient R2, and loss value. For all forecasted data in the validation step, the R2 value was calculated for one year of predicted data. The resulting model uses a logic gate to select the MoE for the specific fisheries species and region, then the data is normalized with a min-max function and reshaped with the delayed input in new columns; this new data frame is passed to the MoE to make the predictions, another output gate pass the values to a denormalization function (Figure 2).

**Figure 2.**
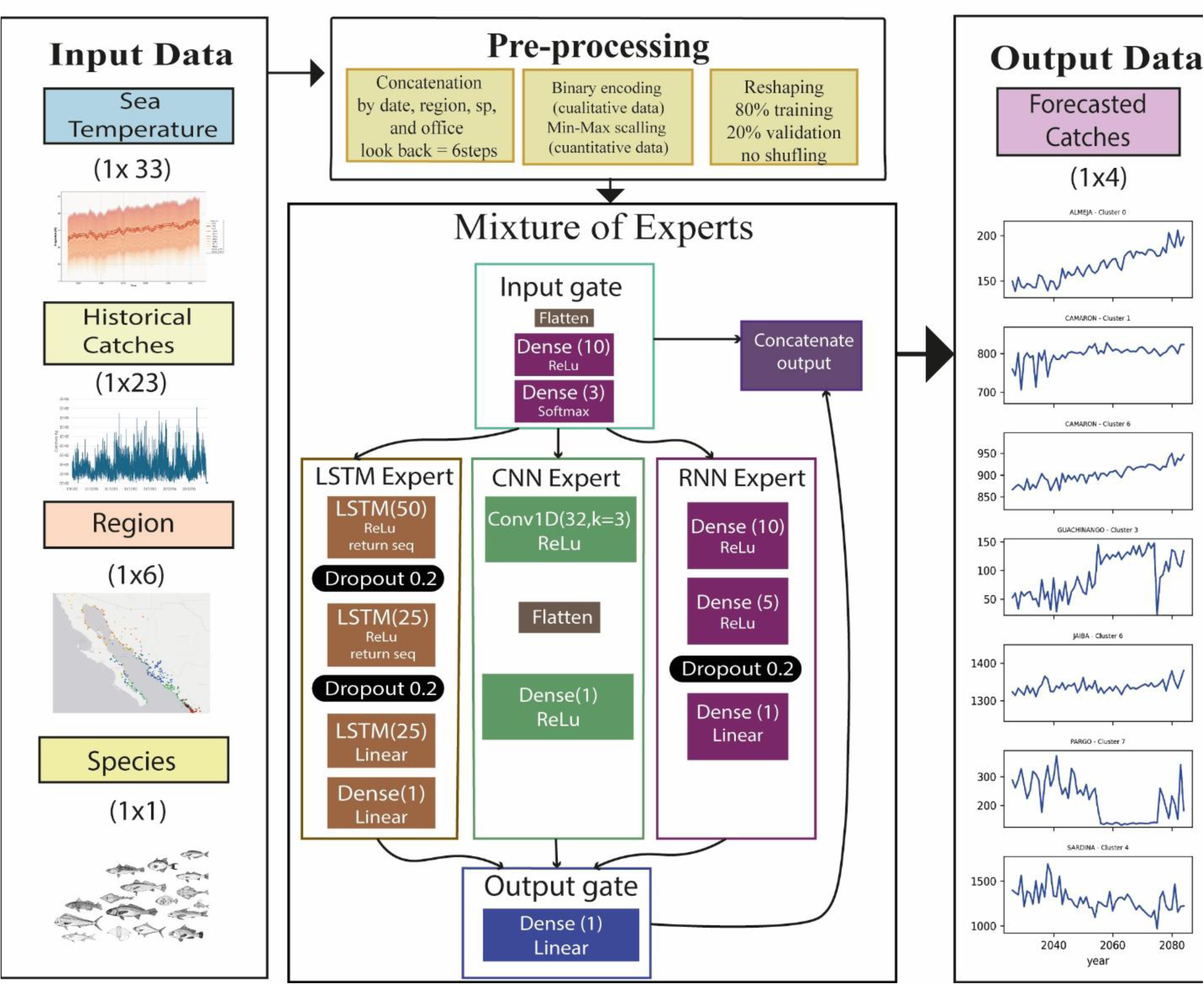
MoE Model.

### Sensitivity analysis

A Shapley Additive Explanations (SHAP) analysis was applied to evaluate the importance of features for each predictive model [73]. SHAP values were computed to quantify the contribution of each feature to the predictions. The aggregated results provided a detailed understanding of the importance of features for each species and fisheries region, enabling robust comparisons across models. For the analysis, we evaluated the SHAP value for each species in the habitat group and calculated the mean value. This analysis uses historical and forecasted data from 2020. This helped identify the main factors determining the forecast and how these change over time. Additionally, we mapped the activation values of the model’s hidden layers to identify the behavioral patterns of the model in response to different magnitudes in the input data and their effect on the model’s output.

A secondary sensitivity analysis explored how the model responds to changes in water temperatures at various depths. We created multiple temperature scenarios ranging from 15°C to 32°C, with increments of 1°C. Each temperature was applied uniformly across all ocean depths. The model predicted biomass (landed_w_kg) for every scenario and we calculated descriptive statistics such as the mean, maximum, minimum, and standard deviation. This process shed light on how variations in water temperature influenced the model’s predictions, offering valuable insights into the system’s thermal sensitivity. It also helped identify critical thresholds or optimal temperature ranges for each species and fisheries region under study.

We also monitored the model’s neuron activation during the forecasting process, and compared it with the input data. This process aims to explore how the model reacts to the input variables.

### Economic analysis

For the economic analysis, we adopted a straightforward approach to quantify the potential economic impact of projected changes in biomass across different fisheries. We first calculated the projected percentage variation in catches for each species group and habitat, based on our predictive models. Subsequently, these percentage changes were applied to the historical average economic value of each fishery, obtained from official catch records and commercial values. This method allowed us to translate projected ecological variations into monetary estimates, expressed in US dollars (using an exchange rate of 20 Mexican pesos per dollar), to evaluate the differential economic impact by region and taxonomic group. While we recognize the limitations of this simplified approach, which does not incorporate price fluctuations or changes in market demand, it provides a valuable first approximation of the economic magnitude of climate change impacts on Gulf of California fisheries.

## Results

We obtained various performance indices, including the mean square error, correlation coefficient R, accuracy, and root mean square error, to assess the model’s performance regarding validation forecasting. We observed (Figure 3) that the correlation coefficient’s performance ranges from 0.54 to 0.98. In contrast, the standard deviation index’s validations ranged between 0.38 and 1.17, indicating the model has forecasting capability and adapts to different scenarios, regions, and fisheries.

**Figure 3.**
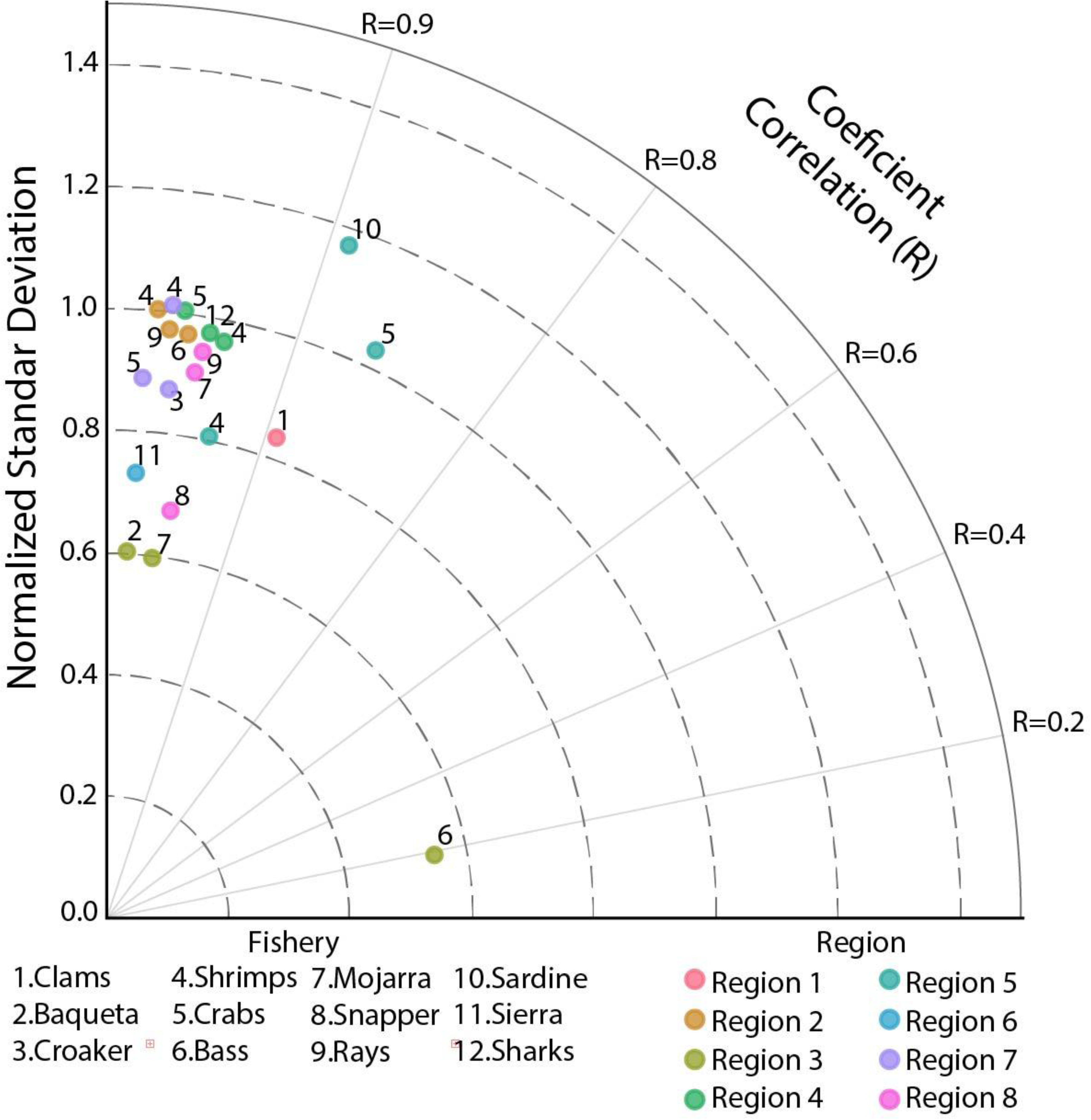
Model performance in validation for different fisheries; the best three validations per region.

The accuracy values exhibit considerable variation contingent upon the specific region and species, oscillating between -207% in Region 6 and 98% in Region 4. The length of the validation sample influences these values, and the specific amount of available historical data in certain fisheries over the regions, for a deeper analysis refer to S5. To contextualize the model’s efficacy, we conducted a comparative analysis of the performance of our Mixture of Experts (MoE) model against the results of models published by other authors. Table 3 provides a contextual comparison between our Mixture of Experts (MoE) model and other relevant deep learning approaches applied to fishery prediction across diverse marine ecosystems. Each study utilized its own regional datasets and specific methodologies, so this comparison should be interpreted from a methodological perspective rather than as a direct performance evaluation under identical conditions. We did not apply these external models to our Gulf of California dataset, but compiled their reported results to contextualize our approach within the current state of the art. Despite differences in underlying data, input variables, and geographic regions, this comparison offers valuable insight into how various deep learning architectures address fishery prediction challenges. Our MoE model achieves competitive performance metrics, with maximum correlation coefficients of 0.98 and accuracy up to 98% under optimal conditions.

**Table 3.**
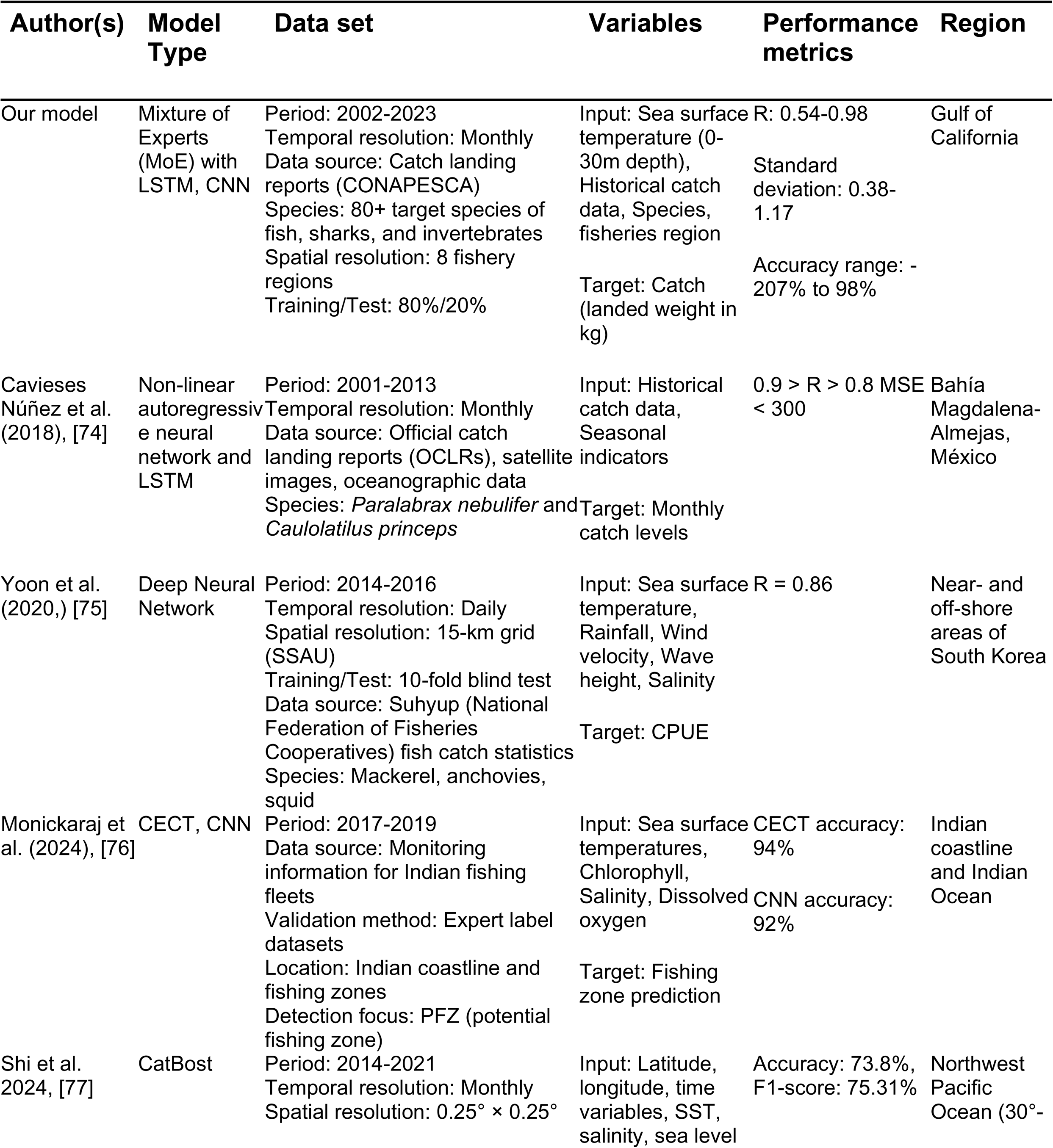
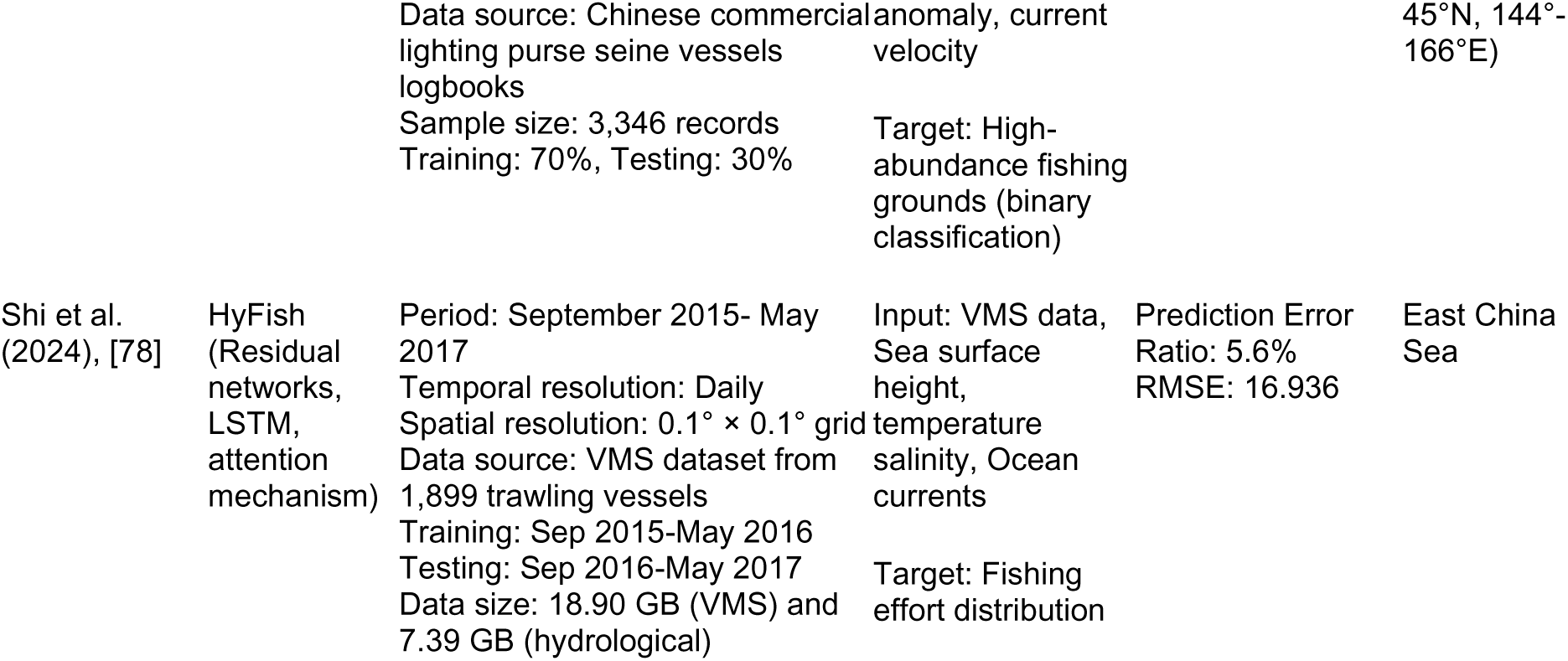
Comparison of models’ performance with similar deep learning approaches.

Figure 4 presents the analysis of catch predictions that reveal distinct patterns across marine species groups and regions. Benthic invertebrates show increasing modal catch values with decreasing variance over time, while benthic fish maintain stable modal trends with significant catch variations between the 2040s and 2060s. Benthopelagic fish display the highest catch volumes, particularly in region 2, with substantial prediction uncertainty. Pelagic fish exhibit notable variability with decreasing trends in region 4 and 5, potentially indicating changes in species distribution or fishing effort. Reef fish maintain moderate values across most regions except for region 8, which shows higher variability and declining trends. Demersal fish predictions in region 5 demonstrate high catches with substantial temporal variation. These patterns suggest complex dynamics in fishing practices and environmental conditions while recognizing that data gaps might result from reporting inconsistencies rather than from the absence of fishing activity.

**Figure 4.**
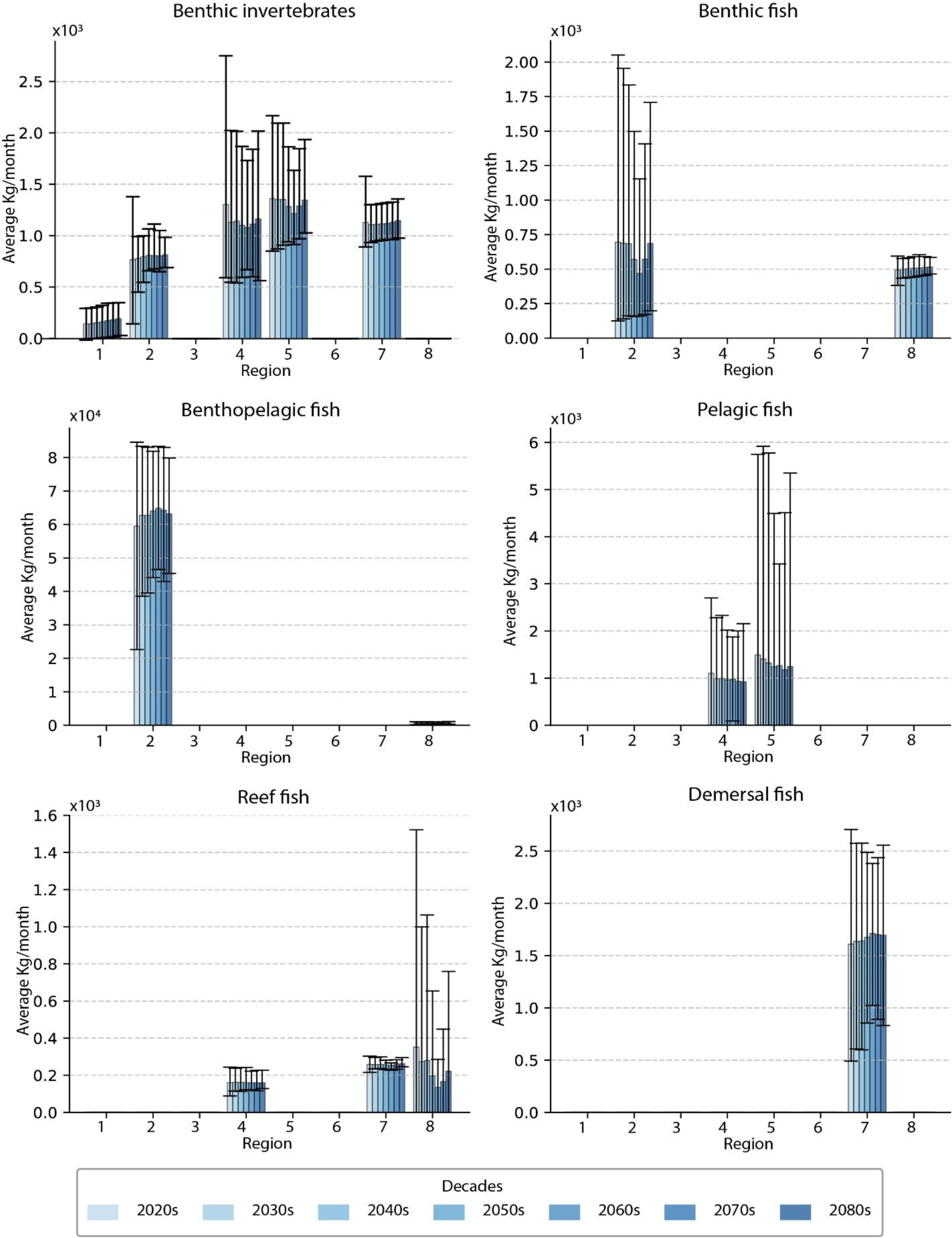
Grouped prediction for different habitats, species, and fisheries regions, minimum and max prediction as whiskers in the plot.

The analysis of projected changes in marine habitat populations across decades (2030s-2080s) in Table 4 reveals diverse temporal patterns. Reef fish and benthic fish display the most pronounced fluctuations, with substantial declines during the 2050s-2060s (approximately -12.46 % and -9.37%, respectively), followed by notable recoveries in the 2070s-2080s (reaching increases of about 10%). Benthic invertebrates show more moderate variations, initially declining by -4.12% in the 2030s before gradually recovering to a 3.18% increase by the 2080s. Benthopelagic fish exhibit an inverse trend, starting with a positive growth of 5.44% in the 2030s but declining to - 1.72% by the 2080s. Demersal fish maintained relatively stable positive values until the 2060s, followed by minimal negative changes. Pelagic fish populations demonstrate considerable volatility, predominantly showing negative changes except for a brief favorable period in the 2060s (1.60%), with variations ranging from -7.90% to 2.67%. These projections suggest significant variability in the response of different marine habitats to future conditions, with some populations showing resilience and recovery potential while others face sustained challenges.

**Table 4.** Variation in the mean forecasted mode of catch by habitat group.

| Habitat group | 2030s | 2040s | 2050s | 2060s | 2070s | 2080s |
| --- | --- | --- | --- | --- | --- | --- |
| Benthic fish | -0.17% | 0.08% | -9.35% | -9.37% | 10.99% | 10.95% |
| Benthic invertebrates | -4.12% | 0.97% | -1.75% | -1.76% | 2.85% | 3.18% |
| Benthopelagic fish | 5.44% | -0.02% | 2.09% | 1.32% | -0.90% | -1.72% |
| Demersal fish | 1.63% | 0.15% | 2.35% | 2.08% | -0.73% | -0.19% |
| Pelagic fish | -7.90% | -3.39% | -4.57% | 1.60% | -5.60% | 2.67% |
| Reef fish | -10.21% | 1.33% | -12.46% | -10.43% | 6.39% | 10.42% |

In contrast, Table 5 presents the economic implications of projected catch changes that vary substantially across habitat groups when applied to historical average values. Reef fish are expected to experience the most significant economic impact, with a projected decline of 1.2 million dollars by the 2050s and a substantial recovery of 1 million dollars by the 2080s. Benthic invertebrates show a concerning trend with decreases of 3.8 million dollars in the 2030s and 1.6 million dollars in the 2050s before showing a remarkable recovery of 2.9 million dollars by the 2080s. In contrast, benthopelagic fish initially showed positive economic returns of 0.5 million dollars in the 2030s, followed by modest gains of 0.2 million dollars in the 2050s and 0.2 million dollars in the 2080s. Demersal fish maintain relatively stable positive economic returns across the decades. In contrast, pelagic fish demonstrate a fluctuating pattern with initial losses -0.9 million in the 2030s, smaller declines (-0.4 million dollars) in the 2050s, and finally showing positive returns (0.3 million dollars) by the 2080s.

**Table 5.** Change in mean forecasted catch value relative to the historical mean value (millions of USD). Exchange rate: 1 dollar = 20 MXN pesos.

| Habitat Group | Average | 2030s | 2050s | 2080s |
| --- | --- | --- | --- | --- |
| Benthic fish | 4.501 | -0.008 | -0.421 | 0.492 |
| Benthic invertebrates | 93.664 | -3.859 | -1.639 | 2.978 |
| Benthopelagic fish | 9.682 | 0.527 | 0.202 | - 0.166 |
| Demersal fish | 13.727 | 0.224 | 0.323 | - 0.0268 |
| Pelagic fish | 12.286 | -0.971 | -0.561 | 0.328 |
| Reef fish | 9.633 | -0.984 | -1.200 | 1.003 |

According to the graphs (Figure 5a and 5b) showing feature importance by habitat using mean SHAP values, distinct patterns can be observed for each group of marine organisms. Benthic fish showed a stronger positive response to temperatures at depths of 18 and 21 m while showing a significant negative reaction at 6 m. For benthic invertebrates, temperatures between 6 and 11 m had an essential positive influence, but a negative effect was observed at 15 m. Benthopelagic fish presented an interesting pattern, with pronounced adverse effects at shallow depths up to 10 m and some positive values at intermediate depths. As for demersal fish, the 6 m depth showed the highest positive importance in the forecast, with a second peak of importance at 13 m.

**Figure 5.**
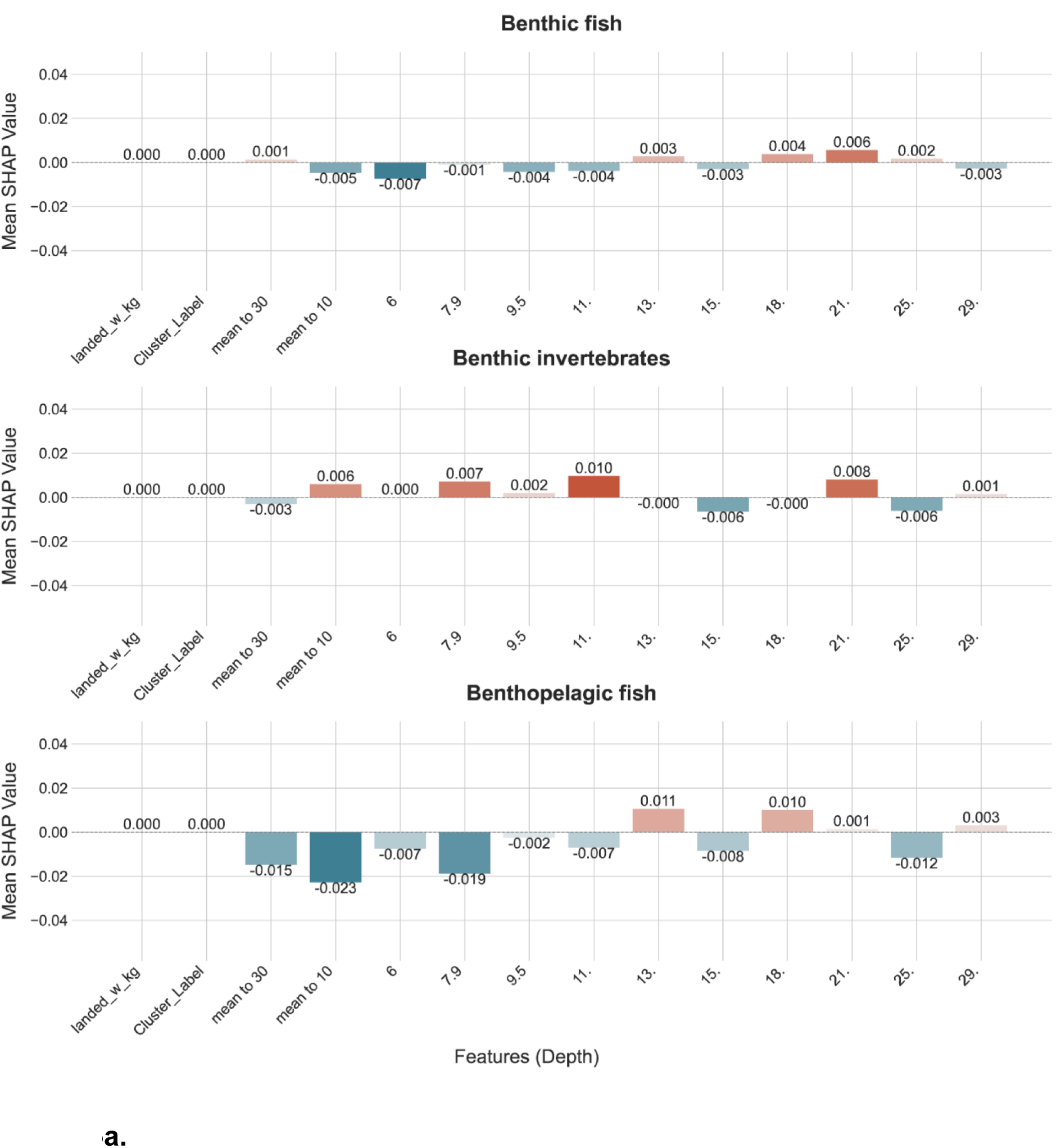
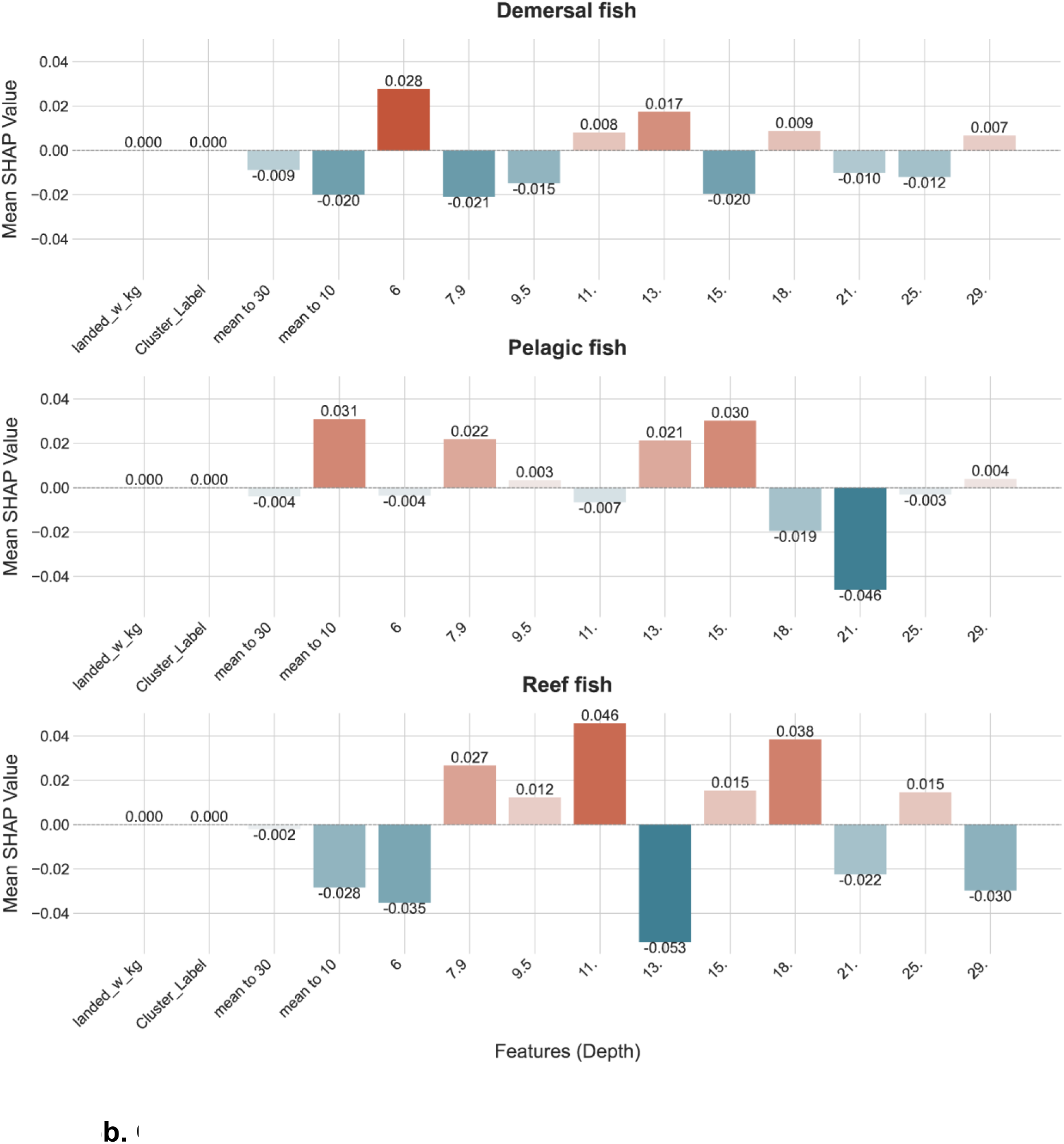
Grouped mean SHAP values for features in the model used to predict landing weight report values for each habitat group.

The temperature prediction model for marine species reveals distinct patterns across different groups, as presented in Figure 6. Most species show significant variability between 15-20°C, with reef fish mainly concentrated between 15-17.5°C. Benthic invertebrates, including shrimp, show peak predictions around 3,000 kg at 15°C, followed by a decline and stabilization at higher temperatures. Other invertebrates (oysters, abalone, clams, and lobster) display gradual increases with temperature, ranging between 1,000 and 2,000 kg. Demersal fish like berrugata and baqueta exhibit high initial predictions (2,000-3,000 kg) at 15°C, while benthopelagic and reef fish species generally maintain predictions below 1,000 kg across temperatures. In Figure 6, Pelagic fish show stable predictions below 1,000 kg, except for Sierra (around 1,500 kg). The model demonstrates prediction convergence as temperatures approach 30-32°C, suggesting common responses to thermal stress, though these values should be interpreted considering biological constraints and model limitations at temperature extremes.

**Figure 6.**
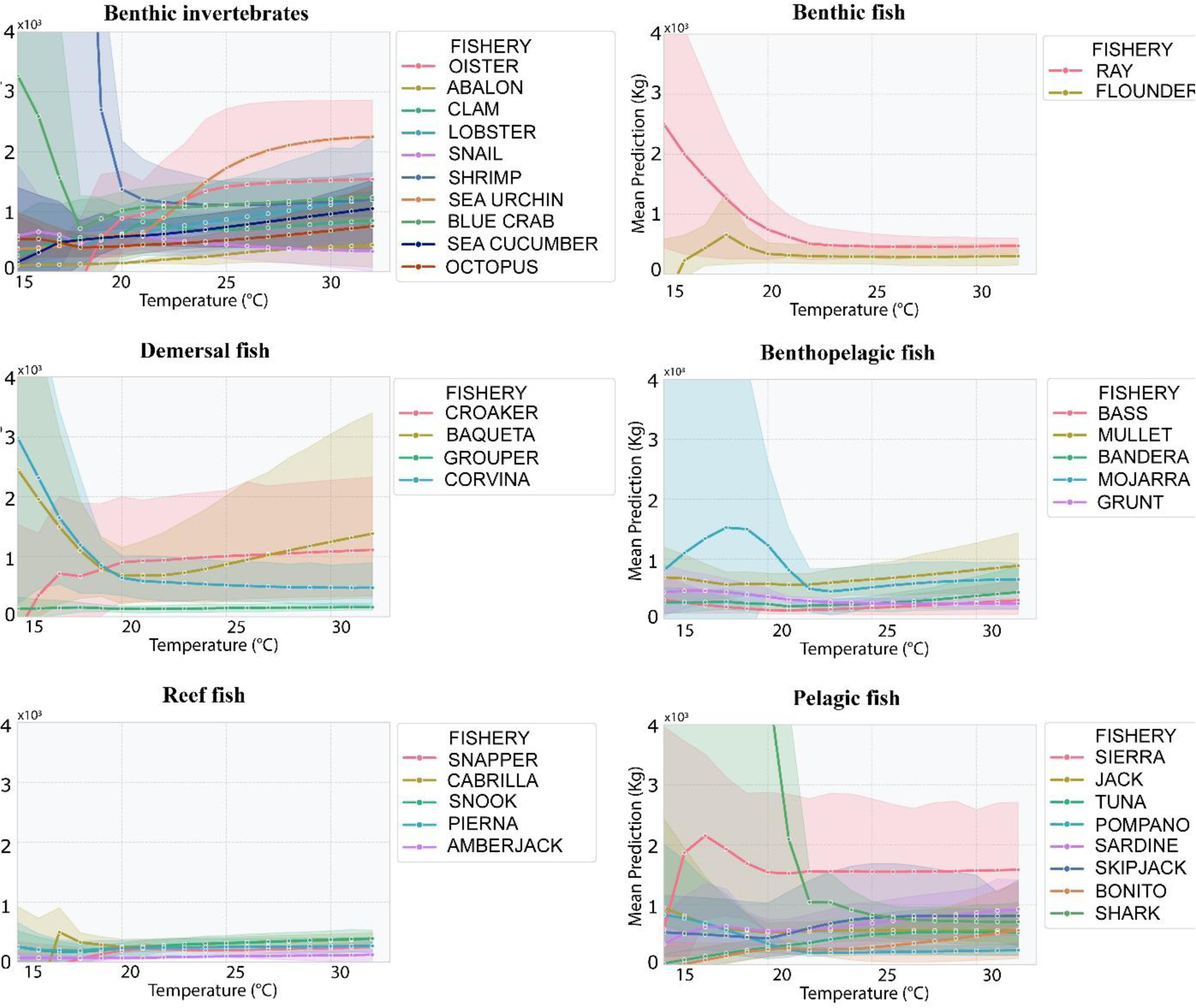
Sensitivity analysis for how temperature impacts the value of landing weight reported for each species group.

The visualization of the MoE (refer to Supplementary Information S3) model when forecasted the corvina catch in region 7 reveals remarkable neuronal specialization: experts with LSTM and CNN structures activate intensely only during high temperature periods starting from 2051, while the RNN expert maintains constant activity throughout the forecast. This functional complementarity demonstrates how CNNs process spatial patterns in thermal profiles, LSTMs capture complex temporal dependencies during extreme events, and RNNs maintain continuity in background signals. For future research, it would be valuable to analyze how the model’s output gate weights these contributions, which could reveal which oceanographic features are determinants for corvina catches under different climate scenarios.

## Discussion

We show how deep learning can be used to predict fish stock behavior. The current ocean warming trend, in combination with large-scale climate variability of the world’s ocean and atmosphere such as the ENSO-events El Niño and La Niña, indicate that we are entering a period that will be characterized by significant changes of SST [27]. In this context, deep learning models will be useful in analyzing the relationship between environmental variables and fishery catch because they can learn from the data about the complexity and non-linearity of these interactions [32].

We highlight significant patterns in projected catches, the sensitivity of marine species to climatic conditions, and the implications for fishery management. Small-scale fisheries’ multispecific and multi-gear characteristics can give them the flexibility to respond to climate change effects and enable them to maximize profitability [31]. Still, sufficient information is needed on future trajectories. Climate change will increase stress, randomness, uncertainty, and disorder in small-scale fisheries, threatening livelihoods [79]. Successful adaptation strategies to climate change in small-scale fisheries, including diversification and adaptive management, will assure good nutrition, food security, and sustainable livelihoods [79].

Our research introduces three significant innovations that differ from previous studies, which have typically focused on a single species or required extensive data inputs. First, we utilize a Mixture of Experts architecture to model nonlinear dynamics in multi-species fisheries, particularly when biological data is scarce. Second, we produce long-term projections for over 80 species across various habitats and regions, extending from the 2030s to the 2080s. Third, we translate projected changes in catch into economic metrics, providing practical guidance for adaptation planning in previously unassessed fisheries. This approach effectively addresses critical knowledge gaps in managing small-scale fisheries in the context of climate change. In this sense, we want to answer the following questions.

The analysis confirmed significant fluctuations in projected catches for the different species groups analyzed between the 2030s and 2080s, followed by a recovery in subsequent years. These results align with previous research documenting the sensitivity of marine invertebrates to changes in temperature and primary productivity, factors that determine their life cycles and distribution patterns [80,81]. This agreement with other authors for some species provides us with a reference point that gives confidence in our forecasts for other species where references are lacking.

### Did the deep learning model demonstrate sufficient predictive efficacy to anticipate future catch volumes for these fisheries despite evolving climate patterns?

The present study employed an MoE model to explore the potential impacts of climate change on small-scale fisheries in the Gulf of California. This model was chosen due to its ability to adapt its internal structure to specialize different parts to specific patterns within the data [82], corresponding to changes in fishery variables related to climate change. The MoE approach was particularly suitable given the nature of small-scale fisheries data, which typically involves multiple species, incomplete information, and varying degrees of data quality across different fisheries [40]. This flexibility was essential for analyzing the vast number of species in the Gulf of California’s small-scale fisheries (more than 80 target species [34]) and addressing the urgent need to provide forecasts to help understand the magnitude of climate change impacts on these fisheries.

When comparing our MoE approach with other deep learning models applied to fisheries (Table 3), a fundamental distinction emerges regarding objective and scope. While previous studies, such as Yoon et al. [75] and Monickaraj et al. [76], have primarily focused on the short-term prediction of high-abundance zones or fishing effort using highly structured datasets, our model addresses the challenge of projecting long-term trends under the SSP5-RCP8.5 climate change scenario using limited and heterogeneous historical data. The multi-specific nature of small-scale fisheries presents a considerably greater computational challenge than the studies conducted by Cavieses Núñez et al. [74] and Shi et al. [77], which focused on only 2 to 5 species. Despite these added complexities, our model demonstrates comparable performance (R: 0.54-0.98) to those more targeted studies. However, our approach’s true strength lies in its predictive accuracy and ability to operationalize results into actionable information for decision-makers. By explicitly modeling the responses of different functional groups across multiple time horizons (from 2030 to 2080) under a high-emissions scenario, we provide a framework for developing region-specific and fishery-specific adaptation strategies, thus facilitating more robust and forward-looking management in the face of potential climate change impacts on highly vulnerable communities dependent on these resources.

Our model demonstrated considerable predictive efficacy during the validation process in forecasting fisheries catch values across various sea temperatures. It is advisable to employ deep learning models for time series forecasting, particularly when biological data is scarce but environmental data is available. The MoE model exhibited strong predictive capabilities for fisheries in the Gulf of California. However, it has certain limitations, especially when forecasted values occasionally fall outside biologically plausible ranges. These challenges often emerged when confronted with environmental conditions not represented in the training dataset, underscoring the difficulties in ecological forecasting in climate change and environmental variability.

The significance of temperature effects was underscored by the SHAP analysis, which identified the most significant depth for forecasting. For example, when comparing benthic fish to reef fish, the SHAP values indicate that temperature strongly influences reef fish. As a novel analytical tool, SHAP has proven invaluable in elucidating the model’s workings [77]. However, while SHAP offers important insights, it cannot fully explain how the model generates its forecasts. Therefore, it is essential to complement this approach with sensitivity analysis and the monitoring of neuronal activation, as this methodology represents a contribution to the implementation of deep learning to fisheries science. By employing this integrated approach, we have gained a clearer understanding of the scenarios that lead to specific model outputs.

### How much will fisheries’ catch be affected under future predictions, and what are the implications of these changes for fishers and fishing communities?

Our model reveals a complex temporal pattern of the impact of ocean warming across the Gulf of California, with significant fluctuations projected between the 2040s and 2060s followed by partial recovery in later decades. Unlike previous studies that focused on single species or uniform regional impacts, our analysis identifies distinct vulnerability profiles across both taxonomic groups and geographic regions.

Most notably, our forecasts predict substantial economic impacts, with benthic invertebrates projected to experience initial losses of $3.8 million in the 2030s followed by a remarkable recovery of $2.9 million by the 2080s. Reef fish show the most dramatic fluctuations, with projected declines of $1.2 million by the 2050s before recovering $1 million by the 2080s. These non-linear responses challenge simpler climate impact models and suggest complex ecosystem reorganizations over time.

The regional heterogeneity in our projections is particularly revealing. We found that 33.3% of regions exhibit a pattern where some species decline while others remain stable, indicating selective vulnerability within local ecosystems. More concerning is the identification of severe vulnerability hotspots such as Region 5, where our model predicts declines across all functional groups, and Region 8, which shows a dramatic 37% decrease in reef fish catches while other species remain relatively stable. This differentiation in climate vulnerability had also been previously linked to changes in spatial distribution for commercial species in the Gulf of California [83]. These patterns align with known thermal responses of key species. The SHAP analysis from our model identified specific depth-temperature relationships that explain these results: benthic fish responded positively to intermediate temperatures (18-21°C), while pelagic fish showed greater stability at lower temperatures. These findings extend previous research on thermal constraints [84,85] by quantifying the economic implications of these biological responses.

Our results complement studies like Cota-Durán et al. [81] on northward shifts in shrimp distribution and Cisneros-Mata et al. [40] on habitat limitation for demersal species, but go further by providing quantitative projections of catch and economic impacts. The fluctuations we observe may be linked to broader climatic patterns, particularly the interaction between ENSO and PDO effects [86][54], but our model provides a more granular view of how these climate drivers translate to specific fishery outcomes across diverse species and regions. The heterogeneity observed in forecasted catch responses to changes in temperature, strongly suggests that climate adaptation strategies must be tailored to specific regions and fisheries rather than applied uniformly across the Gulf. The identification of both vulnerability hotspots and regions of relative stability provides crucial information for prioritizing management interventions and developing targeted support for fishing communities facing the greatest climate risks.

These thermal responses are particularly relevant given that ENSO events suppress upwelling processes, affecting the distribution and abundance of marine species like small pelagic fishes [87]. The projected fluctuations in catch between the 2040s and 2060s identified by our model may be linked to these broader climatic patterns, particularly the interaction between ENSO and PDO effects that have been documented to influence the Gulf’s ecosystem [87,88]. Benthic fish responded positively to intermediate temperatures (18-21°C), while pelagic fish showed stability at lower temperatures. This finding agrees with previous studies showing how thermal constraints influence marine species distribution based on their tolerance ranges. Additional research has identified that extreme temperatures can cause changes in recruitment dynamics and the distribution of key species [80]. For example, it is projected that climate change will cause northward shifts in shrimp distribution in the Gulf of California, directly impacting catches and the local economy [89,90]. Similarly, warming deep waters could limit habitats for benthic fish such as red snapper and lead to distributional changes [83], grouper and baqueta fish may faces similar changes.

Our projections of heterogeneous climate impacts across regions and species align with findings on social vulnerability in fishing communities. Frawley et al. (2019) demonstrated that specialist fishermen, who focus on a limited range of species and often lack ownership of fishing equipment, experience greater vulnerability to environmental fluctuations than generalist fishermen who employ diverse livelihood strategies. These specialist fishers, particularly those dependent on species projected to experience significant range contractions (such as benthic invertebrates in the Gulf of California), may face profound challenges under the climate scenarios we forecast. As Frawley et al. [91] observed, during periods of resource scarcity, specialists often “lacked agency” to modify fishing tactics or switch target species—precisely the adaptive capacity that will be required to navigate the spatially varied impacts our model predicts. This limited flexibility, combined with our projections for some species in critical regions, suggests that climate adaptation strategies must incorporate both ecological forecasts and an understanding of the differential social vulnerabilities within fishing communities.

### Projected catch trajectories will have economic and management implications

The economic implications of our projected catch changes become particularly significant when considered alongside the documented vulnerability of coastal communities in the Gulf [92]. What is highlighted is how these communities’ high dependence on fisheries and limited socioeconomic diversification increase their vulnerability to climate change [92,93]. Our economic projections, which show substantial variations in catch value across different habitat groups, underscore the need for adaptation strategies that consider both ecological changes and socioeconomic impacts. The model’s ability to project these changes across different time horizons could help inform such adaptation planning.

Our analysis reveals significant regional heterogeneity in projected economic impacts (refer to Supplementary Information S4) across different habitat groups along the north-south gradient of the Gulf of California. These economic implications become particularly significant when considered alongside the documented vulnerability of fishing communities in the Gulf, as highlighted by [92] regarding communities’ high dependence on fisheries and limited socioeconomic diversification. Benthic invertebrates, which include economically important species like shrimp, show the most substantial economic impacts. The central regions are projected to experience the largest decreases in value (-15.7 million USD) in Region 2; -6.8 6.8million USD in Region 4). Shrimp fisheries are among the most important in the Gulf of California, contributing up to 57% of national production [94]. Similarly, reef fish fisheries show a consistent economic decline from the central to southern Gulf (-2.0 million USD in central region 2; -2.7 million USD in central-southern region 4; -1.4 million USD in southern region 7), reflecting the vulnerability of reef-associated species to climate change impacts [95].

In contrast, benthopelagic fish show positive economic projections across the Gulf, with the most substantial increases projected for the southernmost region (2.2 million USD in region 8), followed by moderate gains in the central and central-southern regions (0.79 and 0.8 million USD in regions 2 and 4, respectively). This spatial variation aligns with [84] findings on the importance of seasonal and spatial variability in shaping regional fisheries resilience. Small-scale fisheries’ multispecific and multigear character provides flexibility to respond to climate change effects [80].

Pelagic fish present variable projections, with significant economic losses in the central regions (-2.5 million USD in Region 2; -1.6 million USD in Region 4) but potential gains in southern areas. This variability reflects the complex interactions between environmental drivers noted by [89], who demonstrated how conservation policies and environmental changes can lead to unexpected shifts in species composition and ecosystem function. Our economic projections, showing substantial variations in catch value across different habitat groups, underscore the need for adaptation strategies that consider both ecological changes and socioeconomic impacts [96] .

## Conclusion

These spatially-explicit economic projections highlight the need for region-specific management strategies along the Gulf’s latitudinal gradient. The variation in projected impacts suggests that maintaining diverse fishing portfolios across regions could help buffer against climate-driven economic losses in specific fisheries. This aligns with the recommendation by Free et al. [97] and Cisneros-Mata [40] that robust fisheries management reforms can offset climate-induced losses. The model’s ability to project these changes across different time horizons could help inform such adaptation planning. However, as demonstrated by Giron-Nava et al. [80], these projections should be interpreted considering both model uncertainties and the complex dynamics of marine ecosystems under climate change. The findings also support Morzaria-Luna et al. [89] findings that the emphasis on balancing ecological and economic trade-offs in fisheries management decisions.

In this context, while our projections identify potential challenges related to climate change, they also reveal significant opportunities for adaptation and growth. Through collaborative work among the diverse stakeholders involved in the fisheries that sustain the coastal communities of the Gulf of California, and by valuing the wealth of local knowledge and unique strengths of each community, we can develop innovative strategies that not only foster resilience but also open new pathways for sustainable prosperity in the face of these anticipated changes. The effective implementation of these initiatives could generate a future different from what is projected, where communities adapt and thrive, challenging the limitations of any current predictive model. Our prediction results assume that fishing efforts and practices remain similar to historical patterns since we did not explicitly model socio-economic changes. Therefore, sudden shifts in policy or market conditions could cause real outcomes to deviate from these climate-driven forecasts.

## Acknowledgements

This research was part of the “Assessing climate change impacts on small-scale fisheries using deep learning” project funded by the Google AI for Social Good program (Grant # PRGP000002917050) awarded to H. Morzaria-Luna and C.S. Tucker (Carnegie Mellon University). We acknowledge support from a fellowship to S. Buechler from The Pennsylvania State University’s Center for Applications of AI & ML to Industry (AIMI). C.S. Tucker provided comments to an earlier version of the manuscript.

## Credit author statement

Conceptualization: Hem Nalini Morzaria-Luna, Ricardo Cavieses-Nuñez, Qingyuan Lu,, Pratyush Mallick, Soundar Kumara;

Methodology: Ricardo Cavieses-Nuñez, Hem Nalini Morzaria-Luna, Qingyuan Lu, Pratyush Mallick, Soundar Kumara;

Formal analysis: Ricardo Cavieses-Nuñez, Qingyuan Lu, Pratyush Mallick; Investigation: Ricardo Cavieses-Nuñez, Hem Nalini Morzaria-Luna, Soundar Kumara; Resources: Hem Nalini Morzaria-Luna;

Data curation: Ricardo Cavieses-Nuñez, Hem Nalini Morzaria-Luna;

Writing - Original Draft: Ricardo Cavieses-Nuñez, Hem Nalini Morzaria-Luna, Claudia Rebeca Navarrete-Torices, Qingyuan Lu;

Writing - Review & Editing: Ricardo Cavieses-Nuñez, Hem Nalini Morzaria-Luna, Claudia Rebeca Navarrete-Torices, Qingyuan Lu, Pratyush Mallick, Soundar Kumara; Karen Lopez-Olmedo; Gabriela Cruz-Pinón, Stephanie J. Buechler

Visualization: Ricardo Cavieses-Nuñez, Claudia Rebeca Navarrete-Torices; Supervision: Hem Nalini Morzaria-Luna, Soundar Kumara;

Project administration: Hem Nalini Morzaria-Luna;

Funding acquisition: Hem Nalini Morzaria-Luna, Soundar Kumara, Stephanie J. Buechler;

## Data availability

Github repository at: cedointercultural/CC-DLM-MFisheries

Data avalible at: https://zenodo.org/records/15053836

## Supplementary information

**S1.**
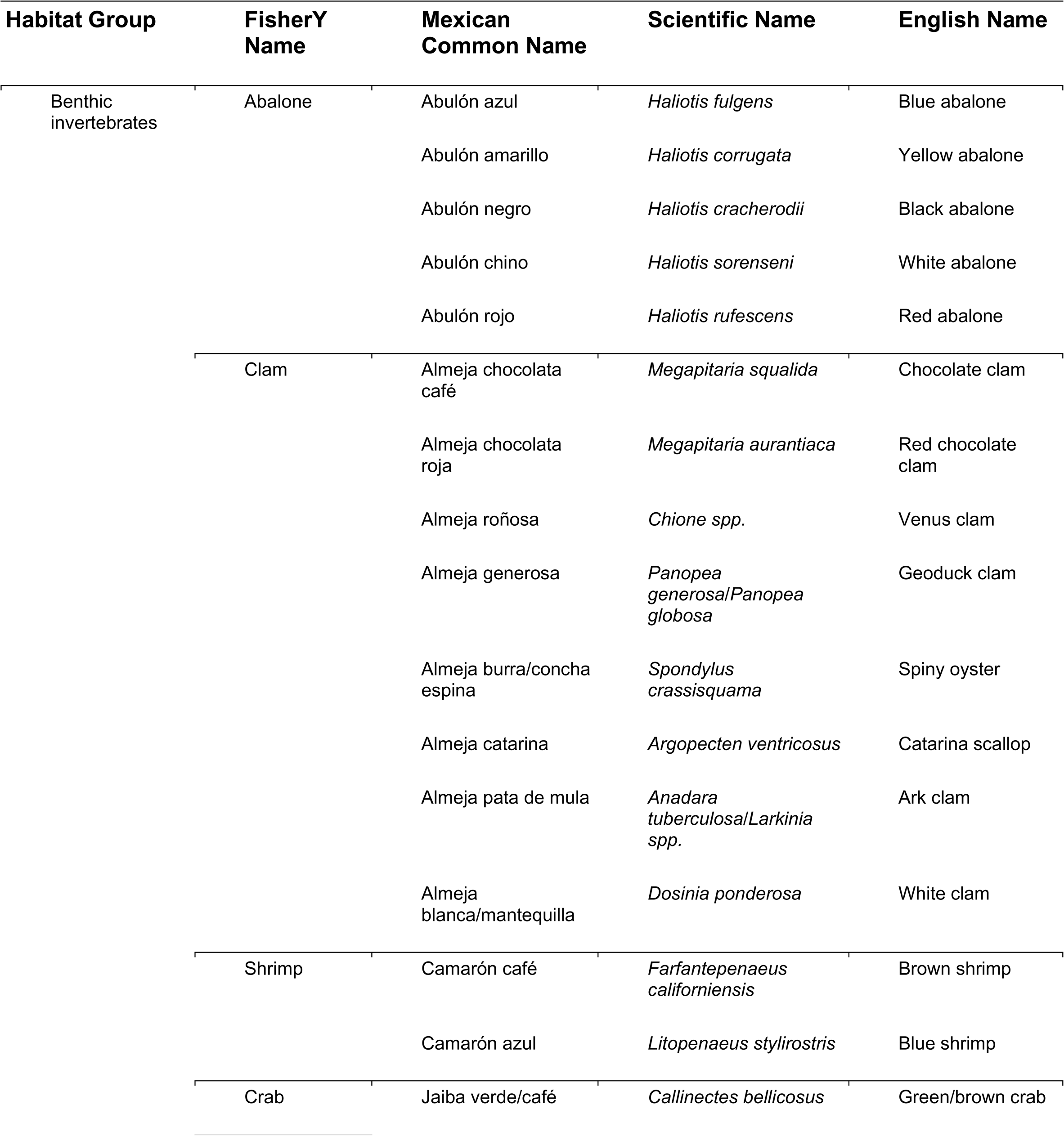
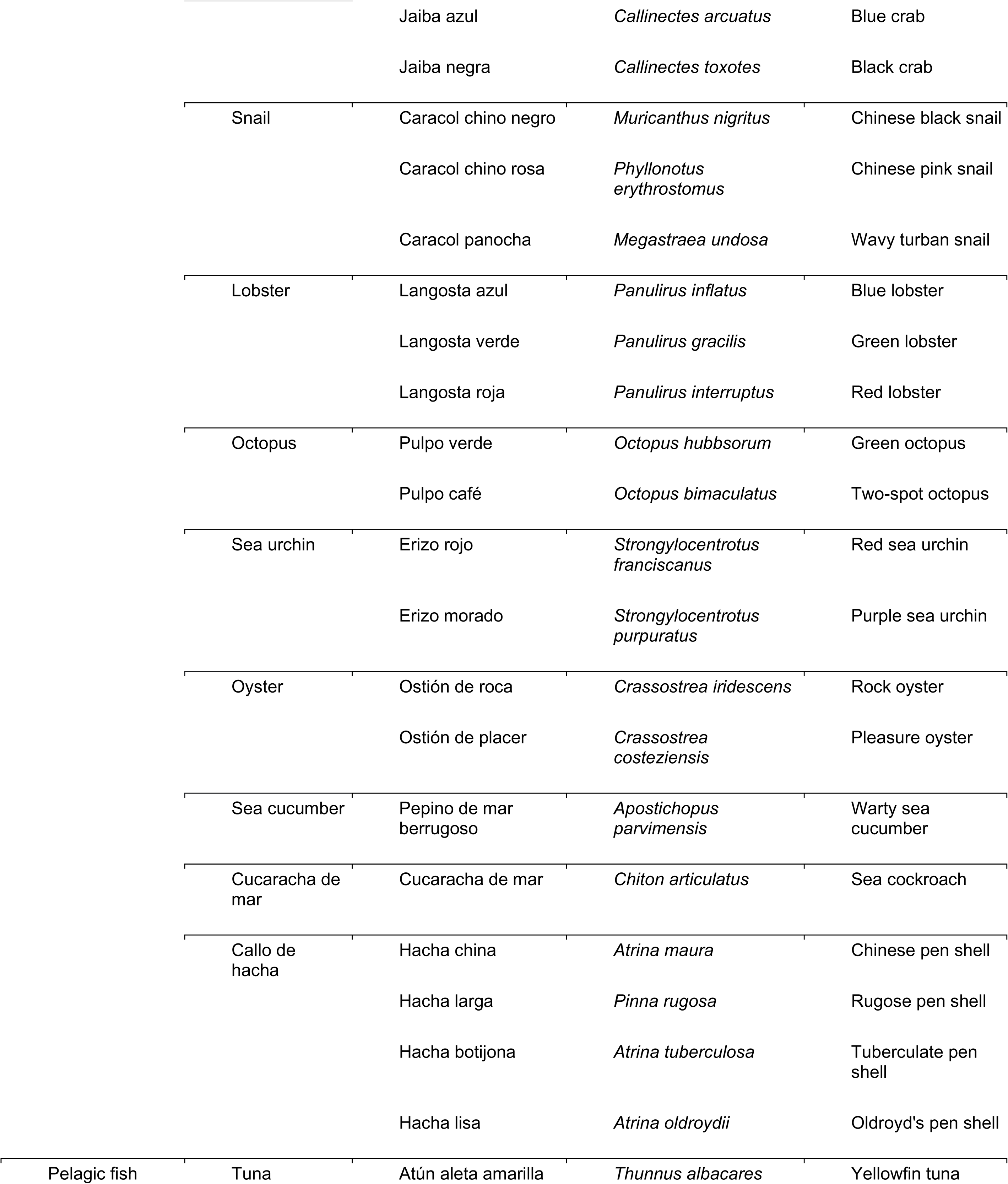
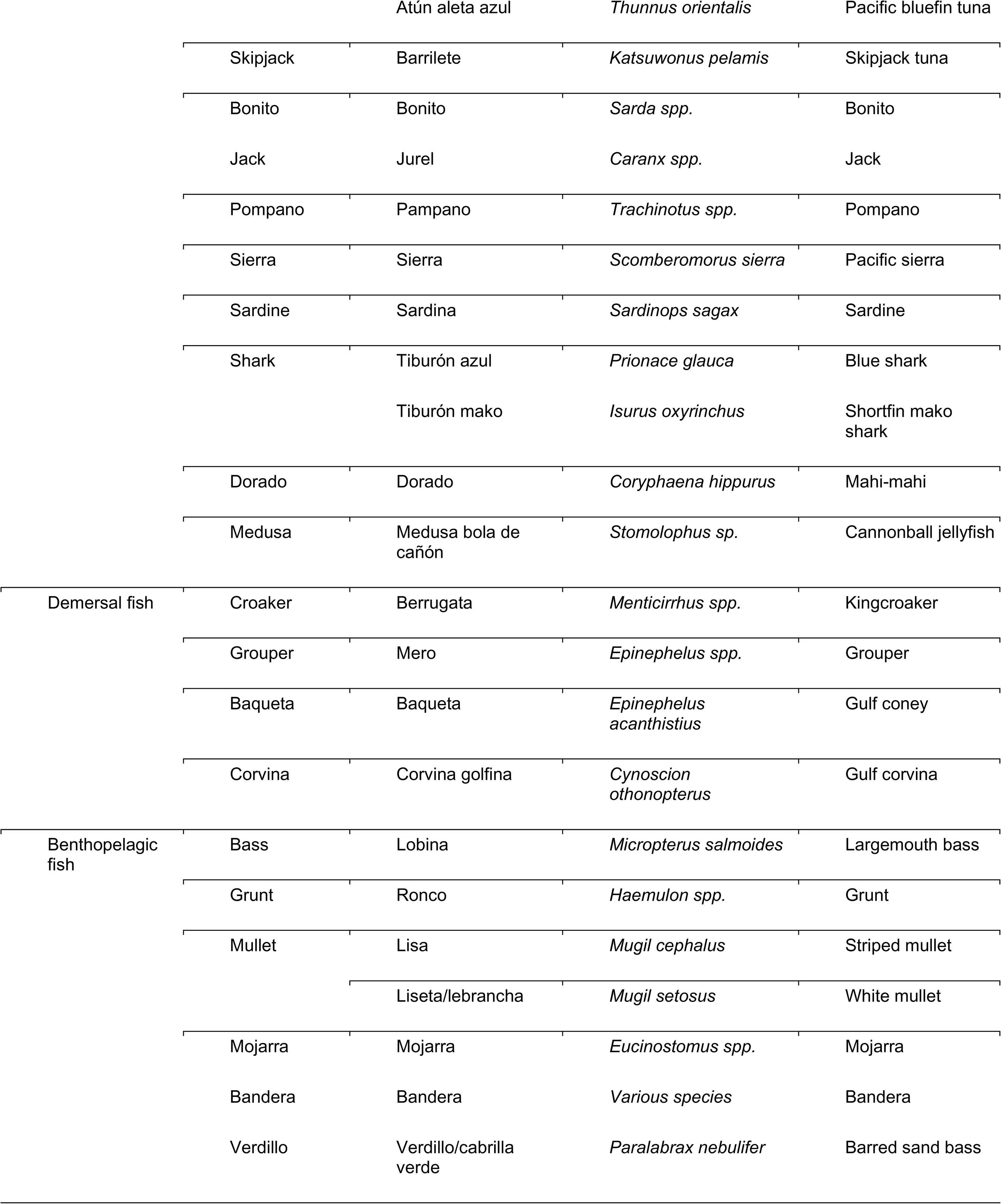
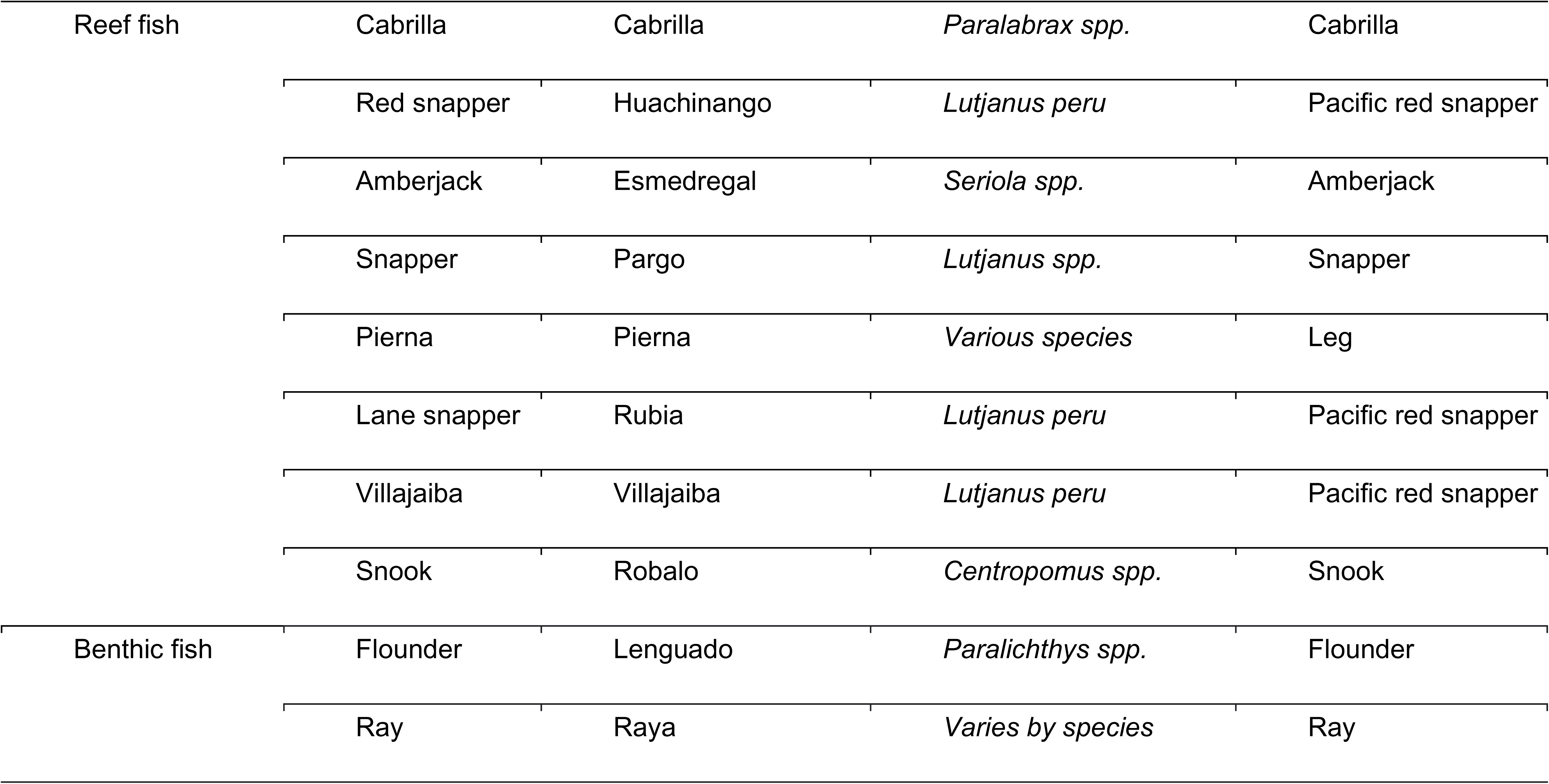
Fisheries Resource Table by Habitat Group.

**S2.**
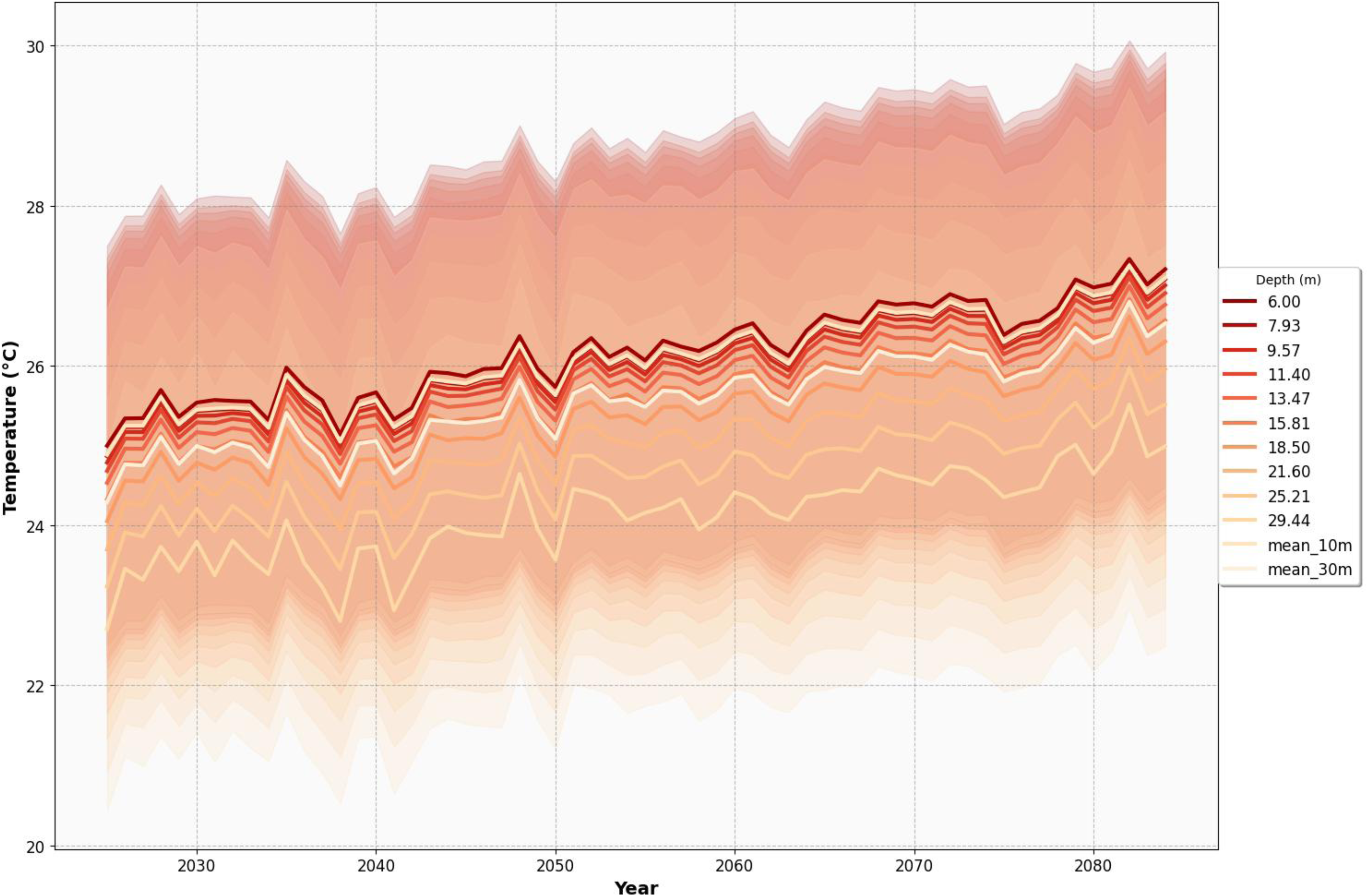
Potential sea temperature with yearly maximum and minimum (colored areas) and the mean temperature in darker lines at a 0.083° × 0.083° resolution for 0-30 m depth. Data from the E.U. Copernicus Marine Service Information Global Ocean Physics Reanalysis https://doi.org/10.48670/moi-00021 from December 1992 to July 2024.

**S3.**
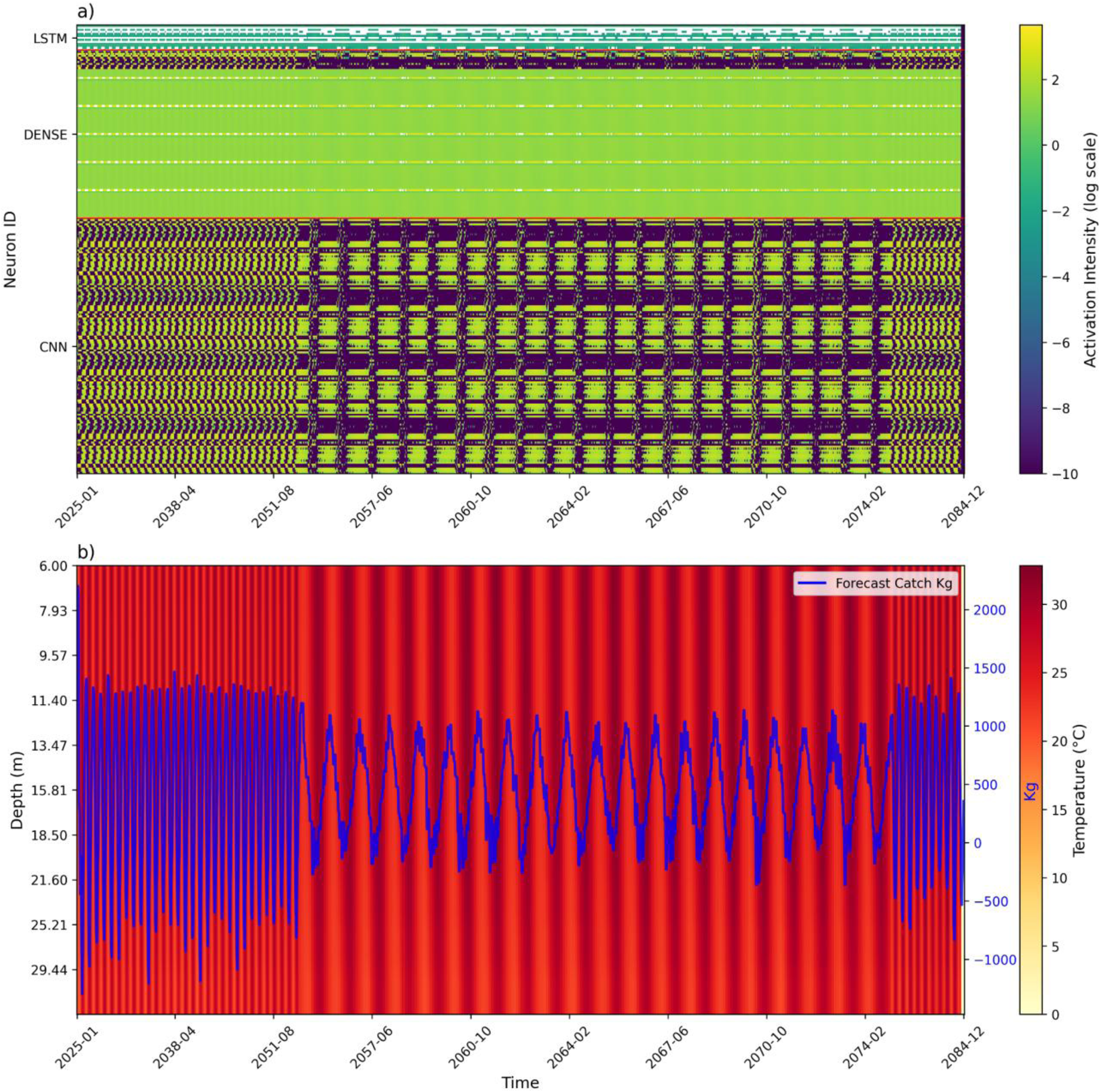
Nodel neuron activation monitoring (a) and forecast for in region six corvina catch response to temperature at different depth (b).

**S4.** Total catches income change predicted for each region.

**S4. Total catches income change predicted for each region.**
|  | Region |  |  |  |  |  |  |  |
| --- | --- | --- | --- | --- | --- | --- | --- | --- |
| Habitat | 0 | 1 | 2 | 3 | 4 | 5 | 6 | 7 |
| Benthic fish | - 0.0007 | -0.02 | -0.0005 | -0.07 | -0.01 | -0.01 | -0.34 | -0.015 |
| Benthic invertebrates | - 0.15 | -15.71 | -0.02 | -6.82 | -0.31 | -0.59 | -5.82 | -1.45 |
| Benthopelagic fish | 0.02 | 0.79 | 0.35 | 0.80 | 0.03 | 0.02 | 0.035 | 2.2 |
| Demersal fish | 0 | 0.355 | 0 | 0.470 | 0.003 | 0.293 | 0.566 | 0.099 |
| Pelagic fish | -0.008 | -2.47 | -0.004 | -1.53 | -0.015 | -0.464 | -2.29 | -0.97 |
| Reef fish | -17.21 | -2,016.69 | -2.34 | -2,705.17 | -12.65 | -5.11 | -1,341.03 | -1,767.98 |

**S5.**
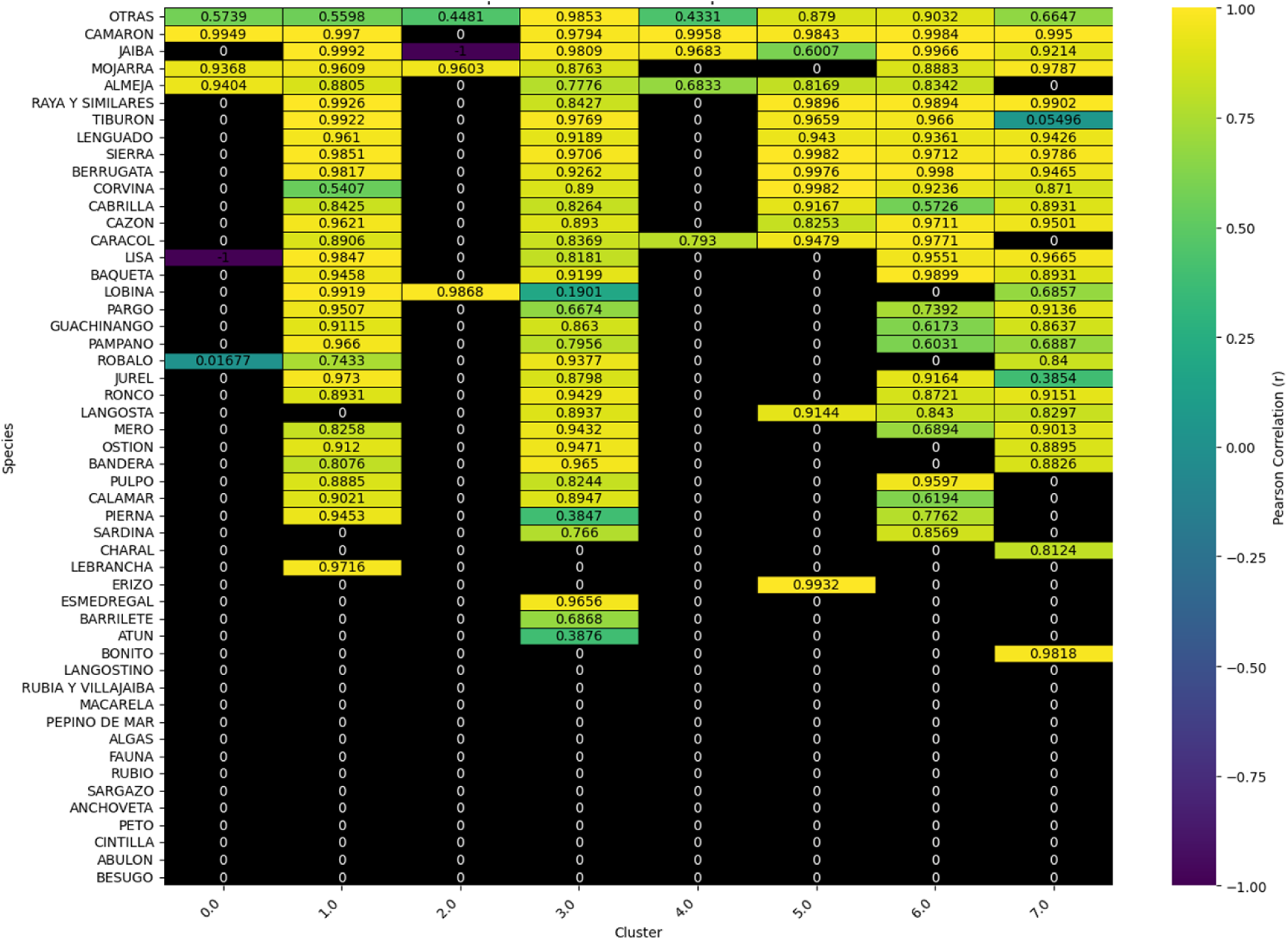
Heatmap of MoE performance in each fisherie for each region.

