## Supplementary figures and images for "Assessing climate change impacts for small-scale fisheries in the Gulf of California using Deep Learning"

### S3

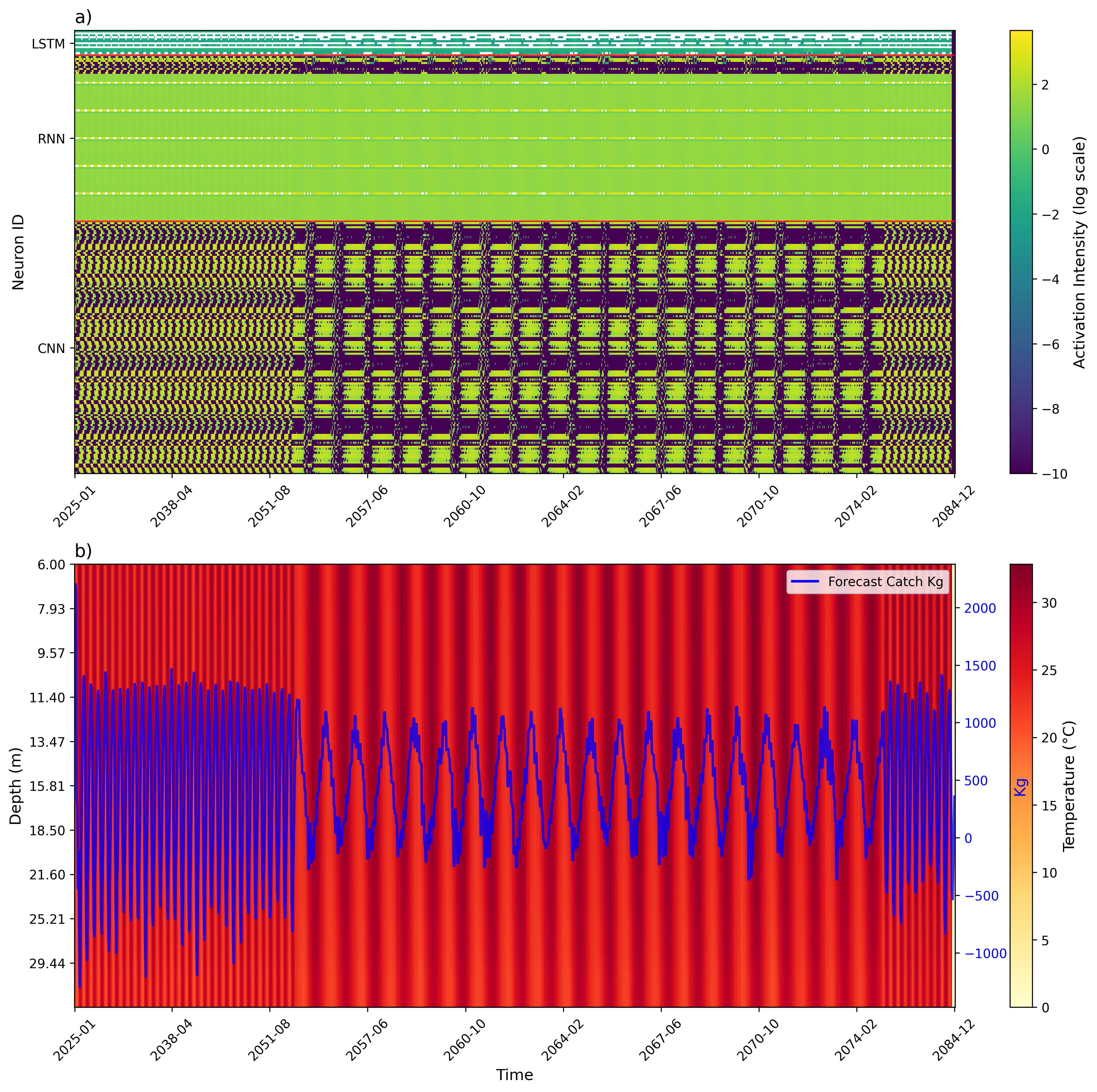
